# Humanized tau and amyloid-β deposition accelerate tau propagation, neuronal cell loss and neurophysiological dysfunction in novel mouse models of primary age-related tauopathy and Alzheimer’s disease

**DOI:** 10.64898/2026.02.23.707081

**Authors:** Arun Reddy Ravula, Ayaka Hagita-Tatsumoto, Brendan J. Gibbs, Nibedita Basu Ray, Justice G. Ellison, Alejandro Longtree-Preciado, Stephanie Radhakishun, Wenhui Qiao, Na Zhao, Takashi Saito, Seiko Ikezu, Tsuneya Ikezu

**Affiliations:** Department of Neuroscience, Mayo Clinic Florida, Jacksonville, Florida, USA; University of Manchester, Manchester, UK; Department of Neurology, Graduate School of Medicine, University of Tokyo, Tokyo, Japan; Mayo Clinic Alzheimer’s Disease Research Center, Mayo Clinic Florida, Jacksonville, Florida, USA

**Author notes:** Correspondence: Tsuneya Ikezu, MD, Ph.D.; Seiko Ikezu, M.D. These authors contributed equally (Arun Reddy Ravula, Ayaka Hagita-Tatsumoto).

**Keywords:** Alzheimer’s Disease, Tau Propagation, Tau Pathology, Braak Staging, Humanized Animal Models, Neuronal Excitability

## Abstract

In Alzheimer’s disease (AD), tau pathology arises in entorhinal cortex layer II (ECII) and advances through defined hippocampal circuits to CA1 and connected neocortical regions, yet the determinants of this hierarchical spread remain unclear. We previously established a circuit-defined propagation model by expressing Cre-inducible human P301L 2N4R tau selectively in Wolframin-1 (Wfs1)^+^ ECII neurons using AAV-FLEX-Tau^P301L^ in Wfs1-Cre mice. Here, to test how amyloid-β (Aβ) and human tau background shape propagation, we generated human *MAPT* knock-in Wfs1 mice and APP^NL-G-F^/*MAPT* double knock-in Wfs1 mice (T-Wfs1 and AT-Wfs1) and induced ECII-restricted Tau^P301L^ expression. Three months after injection, phosphorylated or misfolded tau-positive neurons were enriched in proximal CA1 in Wfs1 and T-Wfs1 mice, resembling primary age-related tauopathy, whereas AT-Wfs1 mice showed preferential accumulation near the CA1/subiculum (Sub) boundary, consistent with an AD-like pattern. In T-Wfs1 and AT-Wfs1 mice, tau spread extended through Sub to neocortical regions, and phosphorylated tau accumulated predominantly in excitatory rather than inhibitory neurons.

Electrophysiological analyses revealed increased spontaneous neuronal firing and impaired GABAergic transmission in the CA1/Sub boundary and neocortical areas in T-Wfs1 and AT-Wfs1 mice, indicative of impaired GABAergic input and enhanced neuronal excitability in these regions. Together, these data indicate that human *MAPT* and Aβ pathology shift the circuit topography of tau propagation and are associated with early network dysfunction, supporting a synergistic interaction that promotes AD-like spread and synaptic imbalance.

## Introduction

In Alzheimer’s disease (AD) brain, tau pathology develops in a hierarchical pattern as classified in Braak stages (Braak, Alafuzoff et al. 2006). Phosphorylated tau appears in the entorhinal cortex layer II (ECII) in Braak stage II and accumulates in pyramidal neurons of the hippocampal cornu ammonis1 (CA1) region. This pathology is also seen in primary age-related tauopathy (PART) and represents benign pathology unassociated with cognitive impairments (Crary, Trojanowski et al. 2014). By Braak stage III, the pathology extends into the adjacent neocortex, reflecting the progressive spread of tau pathology from the medial temporal lobe to neocortical areas and often associated with mild cognitive impairment (limbic stage) (Lowe, Wiste et al. 2018, Therriault, Pascoal et al. 2022, St-Onge, Chapleau et al. 2023, St-Onge, Chapleau et al. 2024). Although tau is heavily implicated in the propagation of AD, amyloid-β (Aβ) proteins also play a key component of AD pathology. Aβ deposition follows a distinct progression from tau, with early accumulation in isocortical regions (Thal phase 1), followed by involvement of limbic structures such as the EC, CA1, and insular cortical regions (Thal phase 2), and later extends to subcortical areas (Thal phase 3) (Thal, Rub et al. 2002). Aβ-induced neuronal hyperexcitability, characterized by increased glutamate release, reduced GABAergic inhibition, and elevated postsynaptic calcium levels (Li, Hong et al. 2009, Wu, Guo et al. 2014, Lam, Sarkis et al. 2020), has also been shown to enhance tau release and facilitate its intercellular transfer (Yamada, Holth et al. 2014, Wu, Hussaini et al. 2016). Furthermore, recent studies have demonstrated that Aβ promotes tau spreading by eliciting neuronal hyperconnectivity as observed through functional MRI studies (Roemer-Cassiano, Wagner et al. 2025). However, it remains unclear whether or by what mechanisms Aβ significantly contributes to the early-stage propagation of tau pathology from ECII to neocortical regions in AD. To demonstrates that amyloid pathology synergistically enhances tau propagation from the ECII to the neocortex, we employed Wfs1-Cre transgenic mice (Wfs1 mice) in combination with the Cre-inducible AAV-FLEX viral vector system to selectively express the human P301L mutant 2N4R tau (Tau^P301L^) in Wfs1^+^ neurons, which colocalize with the calbindin^+^ pyramidal neuronal cluster in ECll region and form the temporoammonic pathway connecting ECII and CA1 regions (Kitamura, Pignatelli et al. 2014). This novel approach led to the accumulation of tau predominantly in the CA1 region of the hippocampus after stereotaxic injection of AAV-FLEX-

Tau^P301L^ in the MECII region (Delpech, Pathak et al. 2021), recapitulating the Braak stages (I–II) of AD and PART. However, tau pathology in this model is mostly limited to tau accumulation in the CA1 region. To recapitulate the next stages of tau pathology, Wfs1 mice, which had knock-ins for both humanized *TAU* and *APP^NL-G-F^* genes, were injected with AAV2/6-FLEX-Tau^P301L^ into the medial ECII (MECII) of for Cre-inducible expression of Tau^P301L^ in Wfs1^+^ cells. To understand how tau moves through specific neural circuitry, tau deposition path along with cell-specific tau accumulation were observed through immunohistochemistry. Additionally, the effects of the endogenous human tau and Aβ on the intrinsic neuronal excitability were evaluated through field recordings, both in the presence and absence of a GABA_A_ receptor antagonist for the assessment of GABAergic input in these regions. Through this integrative approach, we demonstrate that amyloid pathology and human tau expression synergistically heighten neuronal excitability and enhance tau phosphorylation and propagation from the MECII to the hippocampal/neocortical region, and potentially contributing to network dysfunction in early AD.

## Materials and Methods

### Animals

All mouse care and experimental procedures were approved by the Institutional Animal Care and Use Committee at the Mayo Clinic. Mice were caged in accordance with their own sex and housed in a barrier facility with 12 h light and 12 h dark cycles. Food and water were provided ad libitum. Throughout the life of all mice, veterinary staff closely monitored animals for complications. Wfs1-Cre recombinant mice (Wfs1 mice) in C57BL/6 background were obtained from Tonegawa laboratory (The Picower Institute for Learning and Memory, Massachusetts Institute of Technology) (Kitamura, Pignatelli et al. 2014). Human *MAPT* homozygous knock-in mice (Saito, Matsuba et al. 2014) and *APP*^NL-G-F^ homozygous knock-in mice (Saito, Mihira et al. 2019) in C57BL/6 background were obtained from the RIKEN (Drs.

T.C. Saido and T. Satio) and cross-bred with Wfs1 mice to generate *MAPT*:Wfs1-Cre (T-Wfs1) mice and *APP*^NL-G-F/NL-G-F^:*MAPT*:Wfs1-Cre^+/-^ (AT-Wfs1) mice. Both sexes were used for all the studies and genotyping for animals that were performed by TransnetYX.

### Stereotaxic surgeries

For stereotaxic injection we followed similar methods which we have previously published (Delpech, Pathak et al. 2021). Prior to injections, mice were deeply anesthetized using 1 to 2% isoflurane in a 95% O_2_ and 5% CO_2_ mixture. Injections were then performed using an automated mouse brain stereotaxic injector (Neurostar, Germany) equipped with a pulled borosilicate glass capillary needle (Sutter) filled with mineral oil. The glass needle was lowered to the target site at an insertion speed of 0.1 mm/sec. Once the tip reached the target site, solutions were injected at a rate of 200 nL/min. The needle remained at the target site for 5 min after the injection.

For detection of tau propagation path from ECII, AAV2/6-FLEX-Tau^P301L^ (700 nL, 4.43×10^12^ GC/mL, Addgene 176267, Ikezu Laboratory) was unilaterally injected into the right medial entorhinal cortex layer II (MECII) of seven-month-old mice at Bregma coordinates: anteroposterior (AP) –4.85 mm, mediolateral (ML) +3.45 mm, dorsoventral (DV) –3.30 mm from the skull. Animals were euthanized for immunohistochemical analysis three months post injection. For detection of tau propagation path from Sub or CA1, AAV2/6-FLEX-Tau^P301L^ (700 nL, 4.43×10^12^ GC/mL) was unilaterally injected into the right Sub (AP –3.45 mm, ML +2.06 mm, DV –1.80 mm) or CA1 (AP –2.10 mm, ML +1.70 mm, DV –1.40 mm) of Wfs1 mice at seven months of age. Animals were euthanized for immunohistochemical analysis one month post injection.

### Immunofluorescence staining

Animal brain tissue was harvested after transcardial perfusion with 40 mL ice-cold phosphate-buffered saline (PBS) and 4% paraformaldehyde (PFA) in PBS followed by overnight fixation in 4% PFA in PBS at 4 C°. The following day, brain tissue was transferred to cryoprotection solution containing 30% sucrose in PBS and incubated for two days at 4 C° and then embedded in the cryosection mold (Peel-A-Way^®^ Embedding Mold, Truncated-T8, Polysciences, Cat# 18985-1) with O.C.T. compound (Fisher Scientific, Cat# 23-730-571) and frozen at-80°C. The frozen brain tissues were sectioned by cryostat (CRYOSTAR NX50, Thermo Fisher Scientific) at 30 μm into sagittal sections. Sections were washed with PBS for 10 min prior to antigen retrieval in 10 mM tris base/1 mM EDTA in PBS (pH 9.0) at 95°C or in 10 mM sodium citrate (pH 6.0) at 80 C° for 20 min followed by 20 min at room temperature (RT). Sections were washed three times for 5 min in PBS and then blocked in 5% normal goat or donkey serum/5% bovine serum albumin (BSA)/0.3% Triton X-100 in PBS for 1 hour at RT. Primary antibodies diluted with 5% BSA/0.1% Triton X-100 in PBS and sections were incubated overnight at 4°C. The following antibodies and reagents were used for immunofluorescence staining: mouse anti-human tau (HT7; mouse IgG1, 1:1000, Thermo Fisher Scientific, Cat# MN1000), anti-phosphorylated tau at pSer^202^/pThr^205^ (AT8; mouse IgG, 1:100, Thermo Fisher Scientific, Cat# MN1020), anti-misfolded-tau (Alz50; mouse IgG, 1:200, gift from Peter Davies), anti-MAP2 (goat IgG, 1:400, Proteintech, Cat# 67015-1-PBS), anti-Neurogranin (NRGN; rabbit IgG, 1:200, MilliporeSigma, Cat# AB5620), anti-Parvalbumin (PV; rabbit IgG, 1:250, Abcam, Cat# ab181086), anti-Somatostatin (SST; rabbit IgG, 1:200, MilliporeSigma, Cat# SAB4502861), anti-RGS14 (rabbit IgG, 1:4000, Cell Signaling Technology, Cat# 76703), and anti-NeuN (rabbit IgG, 1:1000, Abcam, Cat# ab190565). Sections were then washed three times in PBS and incubated with goat anti-mouse IgG Alexa Fluor™ 488 secondary antibody (1:1000, Invitrogen, Cat# A-11001), donkey anti-mouse IgG Alexa Fluor^®^ 488 secondary antibody (1:1000 Invitrogen, Cat# A-11015), goat anti-mouse IgG Alexa Fluor™ 647 secondary antibody (1:1000, Invitrogen, Cat# A-21235), goat anti-rabbit IgG Alexa Fluor™ 488 secondary antibody (1:1000, Invitrogen, Cat# A-32731), goat anti-rabbit IgG Alexa Fluor™ 647 secondary antibody (1:1000, Invitrogen, Cat# A-21244), for 2 h at room temperature. Following staining, sections were washed with PBS with 0.02% Triton X-100 followed by washing with PBS and then mounted using Fluoromount-G with DAPI (Invitrogen, Cat# 00-4959-52).

### Image processing and quantification

#### Quantification of the signal-positive area occupancy and cell counting

For the analysis of percentage area coverage of HT7^+^, AT8^+^, Alz50^+^, MAP2^+^, NRGN^+^, or counting of PV^+^ cells, large field fluorescence images of the 30-μm thickness sagittal sections of mouse brains with mouse brain atlas coordinates proximal to ML +2.80 and +3.45 mm were acquired using an epifluorescence microscope (Nikon Eclipse Ti, Japan) with Plan Fluor ELWD 20 × Ph1 DM objective at pixel resolution of 1024 × 1024 and lasers at excitation wavelengths of 405, 488, and 647 nm.

Tiled large images of each brain section were taken with DAPI to identify the exact localization of the section and chose the Region of Interests (ROI) for cornu ammonis 1 (CA1), subiculum (Sub), and neocortex (NCx) according to the Mouse Brain Atlas (Franklin July, 2000) (Fig. S1). For the segmentation of CA1-ROI, anterior-posterior boundaries (CA1/CA2 and CA1/Sub boundaries) were delineated based on the shape of pyramidal cell soma, which was clearly visualized by DAPI staining. Additionally, dorsal-ventral boundaries were delineated by the boundary with the corpus callosum (CC) and the dentate gyrus (DG), and then the CA1-ROI was defined to include the stratum lacunosum-moleculare, stratum radiatum, stratum pyramidale, and stratum oriens. For the segmentation of Sub-ROI, the posterior boundaries were delineated by the edge of the CC and the dorsal-ventral boundaries were delineated by the boundary with the corpus callosum (CC) and the dentate gyrus (DG). The segmentation of NCx-ROI was defined as the primary visual cortex (V1) layer 5/6. The anterior, posterior, dorsal, and ventral boundaries were defined, respectively, by an extrapolated line passing through the edge of the dentate gyrus (DG) granule cell layer, the point at which the six-layered cortical lamination was no longer evident, the border with cortical layer IV, and the boundary with the CC (Fig. S1).

Then, percentage of area coverage was calculated as the proportion of the ROI occupied by the signals of interest using ImageJ software. After conversion of an image into 8-bit, binary masks were generated using a defined threshold, and the signal occupancy in each ROI was measured using the function of “Analyze Particles”. The number of PV^+^ cells was manually counted, and cell density (cell counts / mm²) was calculated using ImageJ software.

### Imaging for tracing the circuit between Sub and NCx region

For tracing the circuit of tau propagation, high-resolution Z-stack confocal images of the 30-μm thickness sagittal sections of mouse brains with mouse brain atlas coordinates from ML +1.60 to +3.45 mm were acquired using a Leica SP8 laser-scanning confocal microscope equipped with a 20×/0.8 N.A. objective (Leica, Germany) and lasers at excitation wavelengths of 405 and 647 nm. Confocal stacks in hippocampal/neocortical region were imaged at pixel resolution of 1024 × 1024. Tile images were merged statistically, deconvoluted by Lightning (Leica, Germany), and processed for maximum projection.

### Analysis of the colocalization of AT8 with neuronal-type specific markers

For the colocalization analysis of AT8^+^ cells with neuronal-type specific markers (NRGN, PV, and SST), high-resolution Z-stack confocal images were acquired using a Lightning Leica SP8 laser-scanning confocal microscope equipped with a 20×/0.8 N.A. objective (Leica Lightning, Germany) and lasers at excitation wavelengths of 405, 488, and 647 nm. Confocal stacks in CA1, Sub and NCx region were imaged at pixel resolution of 1024 × 1024 and at voxel resolution of 0.1 × 0.1 × 0.3 μm as described previously (Ruan, Delpech et al. 2020, Clayton, Delpech et al. 2021, Delpech, Pathak et al. 2021).

Colocalization analysis and cell quantification of confocal images were performed using IMARIS software (version 9.8, Bitplane, Oxford Instruments). 3D image stacks were imported into IMARIS, and fluorescence channels were assigned to their respective markers. Cell populations for AT8 and each marker (NRGN, PV, SST) were segmented using intensity-based thresholding and the surface rendering function, and total cell counts were quantified for each marker within the ROI. Colocalization was defined as a minimum of 25% volume overlap between the AT8-positive surface and the corresponding cell-type surface. The total number of colocalized cells and the ratio of colocalization (%) was quantified for each marker.

### Electrophysiological field recordings

Electrophysiological recordings were performed with age-matched non-tg littermates (WT), Wfs1, T-Wfs1 and AT-Wfs1 mice at 7-8 months of age. Each mouse was acutely decapitated, and the brain was rapidly dissected out and mounted in oxygenated ice-cold cutting solution containing 110 mM sucrose, 60 mM NaCl, 3 mM KCl, 1.25 mM NaH_2_PO_4_, 28 mM NaHCO_3_, 0.6 mM sodium ascorbate, 5 mM glucose, 7 mM MgCl_2_ and 0.5 mM CaCl_2_ (pH 7.35) Coronal sections (300-μm thickness) were obtained using a VT-1000 vibratome (Leica) and whole intact sections containing the hippocampus, NCx and CA1/subiculum boundary were immediately transferred to 37**°**C oxygenated artificial cerebrospinal fluid (aCSF) containing 125 mM NaCl, 5 mM KCl, 1.25 mM NaH_2_PO_4_, 15 mM NaHCO_3_, 25 mM glucose, 1 mM MgCl_2_ and 1 mM CaCl_2_ (pH 7.4). Freshly sectioned coronal tissue slices recovered in an oxygenated aCSF bath for 45 min prior to recordings at 37°C. Both sucrose cutting solution and aCSF were made and oxygenated the same day of experimentation. After recovery, a slice was then transferred to a 35-mm dish containing 37**°** C oxygenated to aCSF and placed on an electrophysiological station with aCSF perfusion.

Microelectrodes (4-6 mΩ) were pulled from borosilicate glass (Sutter Instruments) and filled with Ringer solution containing 140 mM NaCl, 5 mM KCl, 1 mM MgCl_2,_ 1 mM CaCl_2_ and 5 mM Hepes (pH 7.4). Electrodes were then lowered to the CA1/subiculum boundary or visual cortex with a three-axis micromanipulator (Sutter Instruments). The tip of the electrode was inserted at a depth of ∼20 μm into the region of interest (ROI), and negative pressure was applied. Once spontaneous spiking was observed, negative pressure was released to restore atmospheric pressure to the recording pipette. Prior to all recordings, tissue health was visually assessed at both high and low magnification to ensure all recordings were from healthy cells. In each coronal tissue slice, spontaneous spiking activity was recorded in a ROI for 10-15 min. For pharmacological experiments involving the GABA_A_ antagonist gabazine (Sigma, Cat# 5059860001), spontaneous spiking activity was recorded in a ROI as previously described for 5-10 min to establish baseline activity. A saturating concentration of gabazine (10 µM) was introduced via a perfusion system, and the tissue was allowed to rest for 5 min before starting another recording. Recordings were compared before and after the introduction of Gabazine to see if any changes in spontaneous spiking activity occurred. Analysis of spontaneous spiking activity was done in Clampfit (Molecular Devices). A threshold analysis (10% of the highest positive peak) was used to determine spontaneous spiking frequency and interspike interval of field recordings. For each 10–15-min acute slice recording, five 1-min periods were randomly selected in the trace for threshold analysis. To assess changes in spiking activity following the introduction of Gabazine, the spontaneous spiking frequency before and after its application was compared to calculate the percent change in spontaneous spiking rate. All spikes were confirmed through a waveform analysis in Clampfit. Average spiking frequency and interspike intervals were then determined for each slice of recording, and groups were compared using One-Way ANOVAs with post hoc Tukey testing. All statistical analysis was done in SigmaPlot (Grafiti).

## Statistical analysis

All data analyses were performed blinded to the genotype or treatment of the animals. Each animal represents an individual biological data point. Replication and sample sizes for all experiments are detailed in the figure legends. Data are presented as means ± SEM. Statistical analyses were performed using Prism (version 10.0, GraphPad). Comparisons between two groups were analyzed for significance using unpaired Student’s t-test after evaluation of the data distribution. Comparisons among three or more groups were analyzed for significance using an ordinary one-way ANOVA with Post Hoc Tukey’s multiple comparison test.

## Results

### Endogenous human tau and Aβ facilitated the propagation of tau and alter the tau propagation pathway

To evaluate of the effect of endogenous human tau and Aβ on tau propagation in Wfs1 mice, distribution of tau was analyzed by immunohistochemistry using HT7 (anti-human tau) antibodies in the brains of animals received stereotaxic injection of AAV2/6-FLEX-Tau^P301L^ in the MECII at seven months of age. (Fig. 1A). Three months after the injection, Wfs1 mice showed preferential HT7^+^ human tau spread in the ipsilateral proximal CA1 pyramidal cells and distal subiculum (Sub) field in hippocampal regions but not in neocortical regions (NCx) at ML +2.80 mm. T-Wfs1 mice showed HT7^+^ human tau spread in the layer 5/6 of NCx as well as proximal CA1 and distal Sub.

**Fig. 1.**
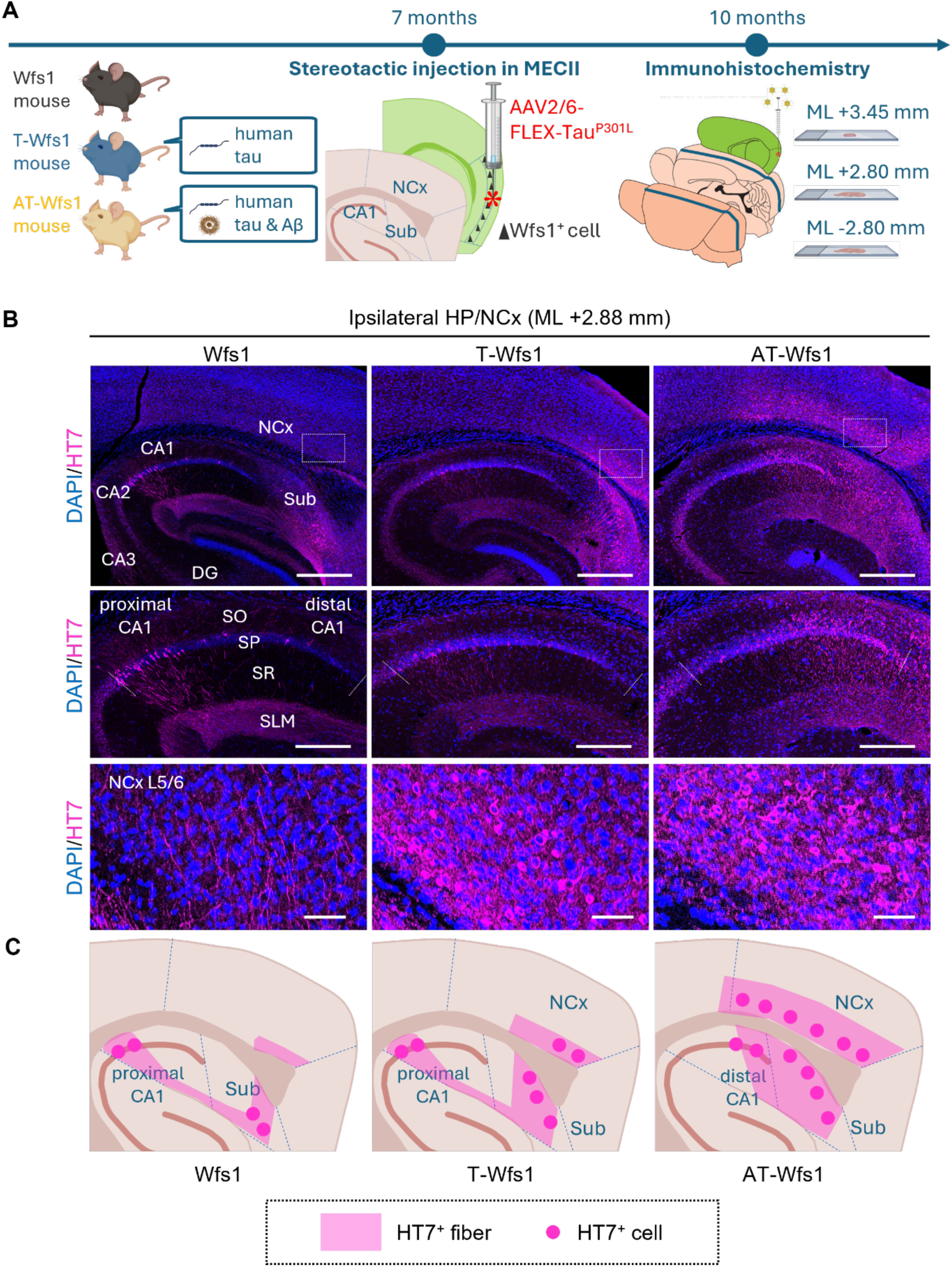
The distribution of propagated tau from MECII to medial hippocampal and cortical region in Wfs1, T-Wfs1, and AT-Wfs1 mice. (**A**) Experimental design schematic for AAV2/6-FLEX-Tau^P301L^ injection in Wfs1-Cre (Wfs1), *MAPT*-KI:Wfs1-Cre (T-Wfs1), and *APP^NL-G-F^*:*MAPT*-KI:Wfs1-Cre (AT-Wfs1) mice. Animals were injected with AAV2/6-FLEX-Tau^P301L^ in the MECII (AP-4.85 mm, ML +3.45 mm, DV - 3.30 mm) at 7 months of age. Mice are euthanized 3 months after the injection for immunohistochemistry. (**B**) **Top**: representative immunofluorescence image of DAPI (blue) and HT7 (total Tau, magenta) in the hippocampal (HP)/neocortical (NCx) at the medial point (ML +2.80 mm). Scale bar = 400 μm. DG: dentate gyrus. **Middle**: Magnified CA1 images. Dotted lines indicate the CA1/CA2 or CA1/Sub boundaries. Scale bar = 200 μm. SO: stratum oriens, SP: stratum pyramidal, SR: stratum radiatum, SLM: stratum lacunosum-moleculare. **Bottom**: Magnified image of boxed area in NCx. Scale bar = 50 μm. (**C**) Summary diagrams of the propagated tau distribution in the three groups.

AT-Wfs1 mice showed HT7^+^ human tau spread widely in the layer 5/6 of NCx as well as distal CA1 and proximal Sub (Fig. 1B, C).

### Endogenous human tau and Aβ accelerated the phosphorylation and misfolding of tau

To evaluate the effect of endogenous human tau and Aβ on tau pathogenesis, the Area occupancy of AT8^+^ p-tau signal and Alz50^+^ misfolded-tau signal were analyzed. In Wfs1 group, both AT8^+^ p-tau and Alz50^+^ misfolded tau were barely detectable in any region at ML +2.80 mm (Fig. 2A) despite the presence of AT8^+^ p-tau accumulation in MECII at ML +3.45 mm (Fig. S3). On the other hand, T-Wfs1 and AT-Wfs1 mice showed strong immunoreactivity of AT8^+^ p-tau and Alz50^+^ misfolded-tau in CA1, Sub, and NCx regions at ML +2.80 mm (Fig. 2A) as well as in the ipsilateral MECII region at ML +3.45 mm (Fig. S3). Interestingly, AT-Wfs1 mice show significant increase in AT8^+^ aera occupancy in the ipsilateral CA1 (1.530 ± 0.3493%, P = 0.0020), Sub (16.72 ± 3.501%, P = 0.0413), and NCx (AT-Wfs1: 16.99 ± 4.937 % vs P = 0.0430) compared to Wfs1 group, whereas no significant difference was observed in T-Wfs1 group in these regions compared to Wfs1 group (CA1: 0.03183± 0.006247%, Sub: 0.6387 ± 0.4347%, NCx: 0.4128 ± 0.08991%; n = 6) (Fig. 2B). In addition, Alz50^+^ area occupancy was significantly increased in the ipsilateral Sub (37.59 ± 5.660%, P = 0.0023) and NCx (25.62 ± 4.322% vs P = 0.0012) of AT-Wfs1 group, but not in T-Wfs1 group, compared to Wfs1 controls (CA1: Sub: 2.926 ± 2.050%, VC: 0.7553 ± 0.2101%; n= 6) (Fig. 2C). Notably, both AT8^+^ and Alz50^+^ signals were negligible in AT-Wfs1 mice injected with saline instead of AAV-FLEX-Tau^P301L^ in the ECII region (Fig. S4), indicating that neither phosphorylation nor misfolding of tau was accumulated by saline injection despite the presence of endogenous human *MAPT/APP*^NL-G-F^ expression.

**Fig. 2.**
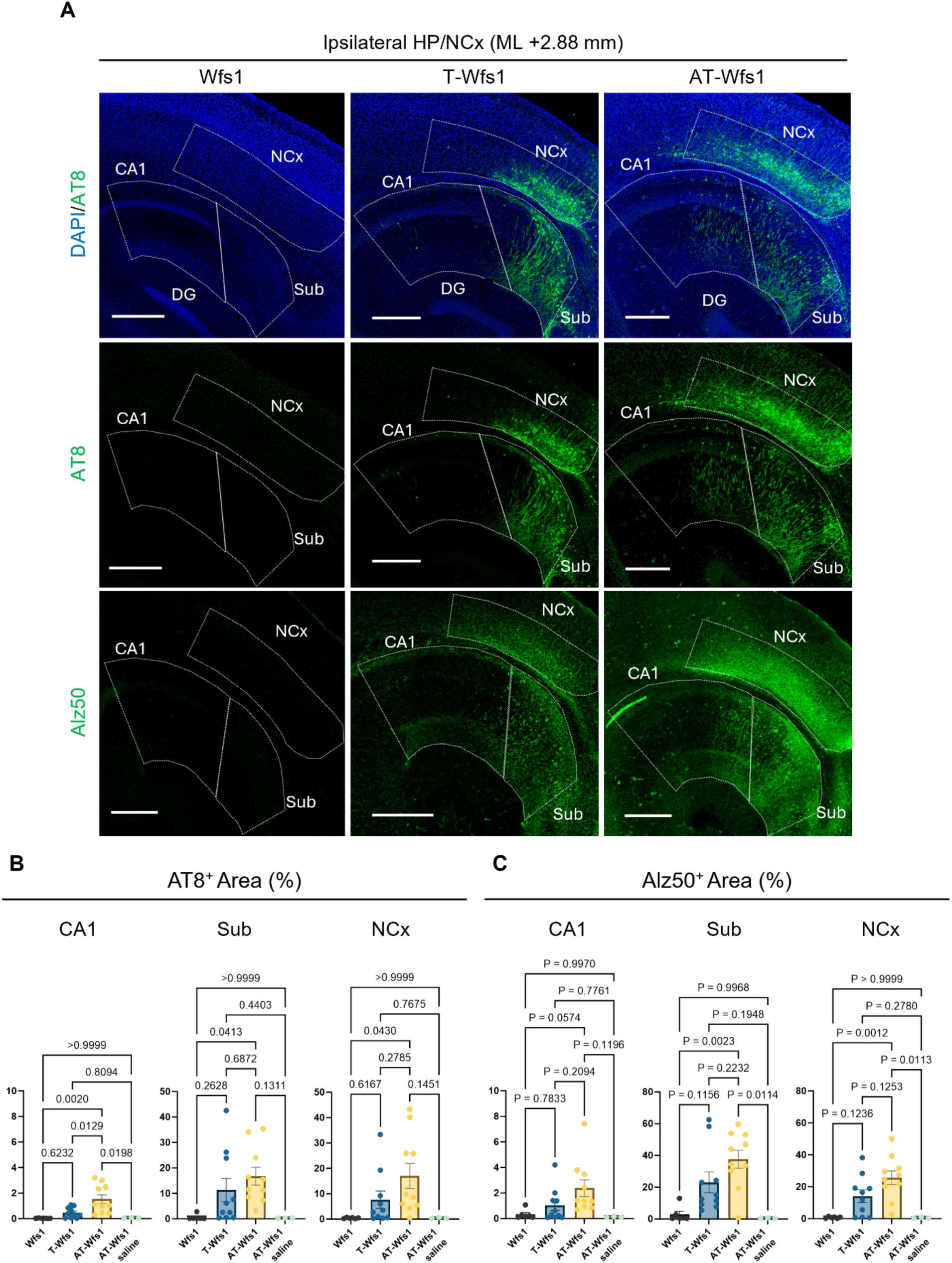
The quantification of p-tau and misfolded tau in the medial HP/NCx region of Wfs1, T-Wfs1, and AT-Wfs1 mice. (**A**) Representative immunofluorescence image of DAPI (blue) and AT8 (pS^202^/pT^205^ Tau, green), and Alz50 (misfolded Tau, green) in the ipsilateral HP/NCx region at the medial point (ML +2.80 mm). Scale bar =400 μm. (**B, C**) Quantification of the AT8^+^ area occupancy (**B**) and Alz50^+^ area occupancy (**C**) in the ipsilateral CA1, Sub, and NCx at ML +2.80 mm. Means ± SEM, Wfs1 n = 6; T-Wfs1 n = 10; AT-Wfs1 n = 10; saline injected AT-Wfs1 (negative control) n = 3 mice. Statistical analyses were performed by one-way ANOVA with Post Hoc Tukey’s multiple comparison test.

In the contralateral hemisphere at ML –2.80 mm, both AT8^+^ and Alz50^+^ cell soma staining were negligible in three groups (Fig. S5B, C), indicate that p-tau and misfolded-tau did not propagate into contralateral hemisphere. However, AT8^+^ and Alz50^+^ puncta were detected mainly in CA3, Sub and NCx of AT-Wfs1 mice. These spots were colocalized with FSB^+^ Aβ plaque in AT-Wfs1 mice (Fig. S5D, E), indicative of plaque-associated dystrophic neurites. Furthermore, in the mice with an extended post-AAV-FLEX-Tau^P301L^ injection incubation period of 6 months, an expansion of misfolded tau propagation toward the anterior cortex beyond V1 region was observed in T-Wfs1 mouse, and an expansion of both AT8^+^ p-tau and Alz50^+^ misfolded-tau pathology toward the anterior cortex beyond V1/V2L/S2 region in AT-Wfs1 mouse (Fig. S6A-C). These results suggest that endogenous expressions of human *MAPT* and APP^NL-G-F^ synergistically accelerate the spread of pathogenic tau to the hippocampal and cortical regions following expression of Tau^P301L^ in Wfs1^+^ neurons in MECII of AT-Wfs1 mice, suggesting the importance of Aβ-related pathology for accelerating the tau propagation beyond hippocampal region.

### The accumulated tau in the CA1/Sub boundary propagated to the deep layers of the adjacent NCx

In T-Wfs1 and AT-Wfs1 mice injected with AAV2/6-FLEX-Tau^P301L^ in the MECII, tau spread into the NCx layer 5/6 which were closely localized to the Sub region where tau accumulated (Fig. 2A). Furthermore, the correlation analysis and liner regression analysis revealed that AT8^+^/Alz50^+^ area occupancy in NCx was positively correlated with that in Sub region (AT8: r = 0.2425, P = 0.0274, n = 20; Alz50: r = 0.2150, P = 0.0395, n = 20), whereas no correlation was observed between NCx and CA1 region (AT8: r = 0.1933, P = 0.0524, n = 20; Alz50: r = 0.03703, P = 0.4163, n = 20) (Fig. S7A, B). We thus hypothesized that the tau propagation to the NCx region is driven by projections from Sub rather than CA1. To address this hypothesis, AAV2/6-FLEX-Tau^P301L^ was injected into either the Sub or CA1 of Wfs1 mice at seven months of age, and the spread of human tau was analyzed by immunohistochemistry using HT7 antibody at one-month post injection (Fig. 3A, B). This is based on the presence of Wfs1^+^ neurons in these regions (Fig. S8a,b). Immunostaining of HT7^+^ human tau in the CA1-injected group showed HT7^+^ tau propagated to both lateral-medial direction in the proximal CA1 but did not propagate to either MEC, Sub or NCx regions (Fig. 3C).Strikingly, Sub-injected group showed a distinct pattern of HT7^+^ human tau signal extending proximal Sub, distal CA1 and the deep layer of NCx regions adjacent proximal Sub (Fig. 3D). These results suggested that the accumulation of tau in the Sub/CA1 boundary region contributed to the spread of tau into the NCx regions.

**Fig. 3.**
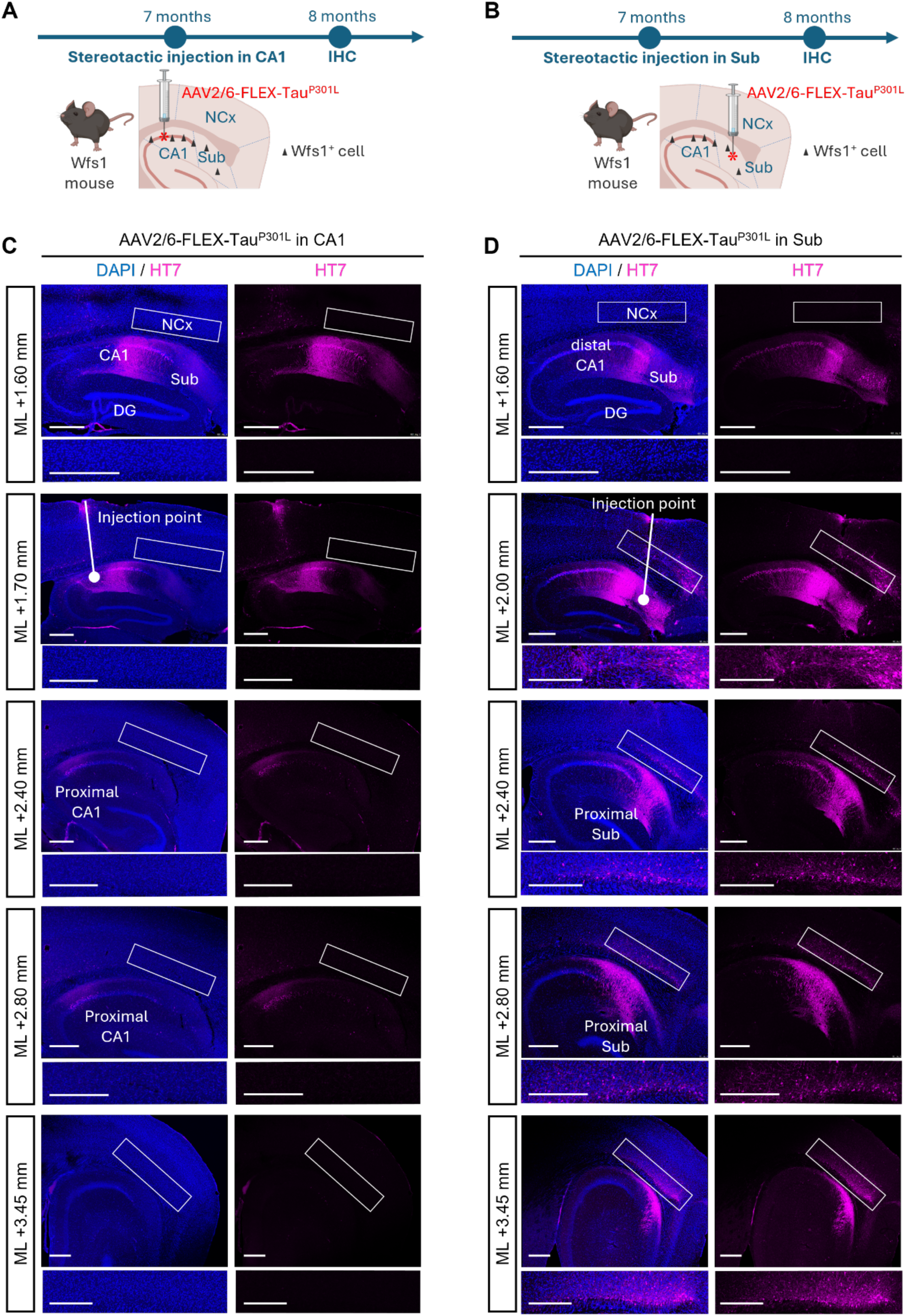
The distribution of the propagated tau in Wfs1 mice injected with AAV-FLEX-Tau^P301L^ in CA1 or Sub (**A, B**) The scheme of the experimental design for AAV2/6-FLEX-Tau^P301L^ injection in Wfs1 mice. Animals were injected with AAV2/6-FLEX-Tau^P301L^ in the Sub (AP-3.45 mm, ML: +2.06 mm, DV:-1.80 mm, **A**) or CA1 (AP-2.10 mm, ML +1.70 mm, DV-1.40 mm, **B**) at 7 months of age. Mice are euthanized 1 month after the injection for immunohistochemistry. (**C, D**) Representative immunofluorescence image of DAPI (blue) and HT7 (magenta) in the sagittal section in ML +1.60, 1.70, 2.00, 2.40, 2.80, and 3.45 mm. The upper panels are hippocampal and cortical region images, and the lower panels are magnified visual cortical region images. Scale bar = 400 μm.

### Tau propagation-independent dendritic loss in the proximal CA1 SR region of AT-Wfs1 mice

We next investigated the impact of dendritic structure in the CA1 stratum radiatum (SR) region for the changes in tau propagation pattern. Apical neuropil atrophy was reported in the CA1-SR region of mild AD cases (Kerchner, Hess et al. 2010). This is dependent on the Aβ deposition as MAP2 reduction in CA1-SR region is correlated with accumulation of Aβ42 in Tg2576 APP Swedish mutant transgenic mice (Takahashi, Capetillo-Zarate et al. 2013). To evaluate of the effect of endogenous human tau and Aβ on the apical dendrite in the SR layer in the proximal CA1, the sagittal sections of contralateral hemisphere were analyzed by immunohistochemistry using MAP2 antibody. MAP2^+^ signal occupancy was significantly decreased in AT-Wfs1 mice, but not in T-Wfs1 mice (Fig. 4A). This result suggested that the dendritic loss in the SR region of proximal CA1 reduces tau targeting proximal CA1 pyramidal neurons, leading to a relative increase in tau accumulation in distal CA1 and the subiculum, followed by enhanced tau spread to neocortical (NCx) regions in AT-Wfs1 mice (Fig. 4B).

**Fig. 4.**
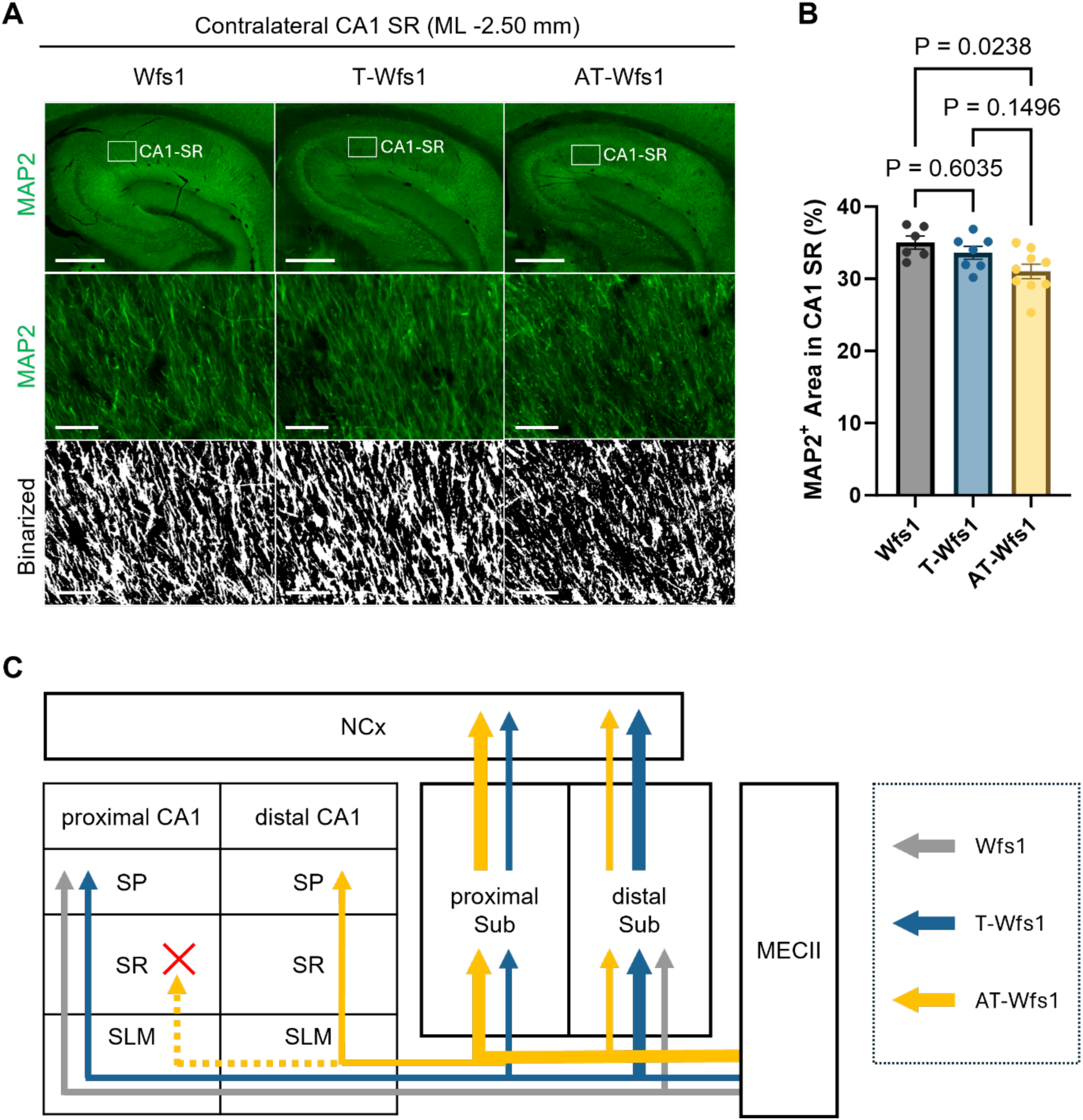
The reduction of dendrite in CA1 SR region of AT-Wfs1 mice. (A) Representative immunofluorescence image of MAP2 (green) in the stratum radiatum (SR) field in the proximal CA1 of the three groups at the medial point (ML-2.80 mm). Scale bar = 40 μm. (B) Quantification of the MAP2^+^ area occupancy in the contralateral CA1, Sub, and NCx at ML +2.80 mm. Means ± SEM, Wfs1 n = 6; T-Wfs1 n = 6; AT-Wfs1 n = 10 mice. Statistical analyses were performed by one-way ANOVA with Post Hoc Tukey’s multiple comparison test.

### Tau propagation-independent impaired GABAergic transmission and promoted neuronal hyper excitability in T-Wfs1 and AT-Wfs1 mice

To investigate the impact of endogenous human tau and Aβ on the baseline of neuronal excitation, field recordings from the CA1/Sub boundary and NCx were conducted for all three groups (Wfs1, T-Wfs and AT-Wfs1 mice, n=5 individuals per group) (Fig. 5 A). Prior to this study, we have confirmed that transgenic expression of Cre in Wfs1^+^ neurons has no effect on neuronal excitability in CA1/Sub and NCx regions compared to non-tg WT littermates (Fig. S9). At the CA1/Sub boundary, T-Wfs1 (0.99 ± 0.25 Hz, P = 0.0006; n=9 slices) and AT-Wfs1 (1.28 ± 0.33 Hz, P < 0.0001; n=13 slices) mice showed significantly higher spontaneous spiking frequencies compared to Wfs1 controls (0.47 ± 0.19 Hz; n=11 slices) (Fig. 5B-D). Interspike intervals were also shorter in T-Wfs1 (996 ± 201 ms, P = 0.0006) and AT-Wfs1 (770 ± 185 ms, P < 0.0001) mice compared to Wfs1 controls (2910 ± 1075 ms) (Fig. 5E). In the NCx, T-Wfs1 (1.17 ± 0.21 Hz, P = 0.0223; n = 8 slices) and AT-Wfs1 (1.74 ± 0.38 Hz, P < 0.0001; n = 10 slices) mice exhibited higher spontaneous spiking frequencies compared to Wfs1 controls (0.79 ± 0.11 Hz; n = 10 slices) (Fig. 5G-I). Interspike intervals were shorter in T-Wfs1 (882 ± 132 ms, P < 0.0001) and AT-Wfs1 (587 ± 119 ms, P < 0.0001) mice compared to Wfs1 controls (1361 ± 210 ms) (Fig. 5J). These findings highlight significantly increased neuronal excitability in both the CA1/Sub boundary and NCx in T-Wfs1 and AT-Wfs1 mice compared to the Wfs1 mice, and the expression of human *MAPT* and *APP*^NL-G-F^ show synergistic effect for the neuronal hyper-excitability.

**Fig. 5.**
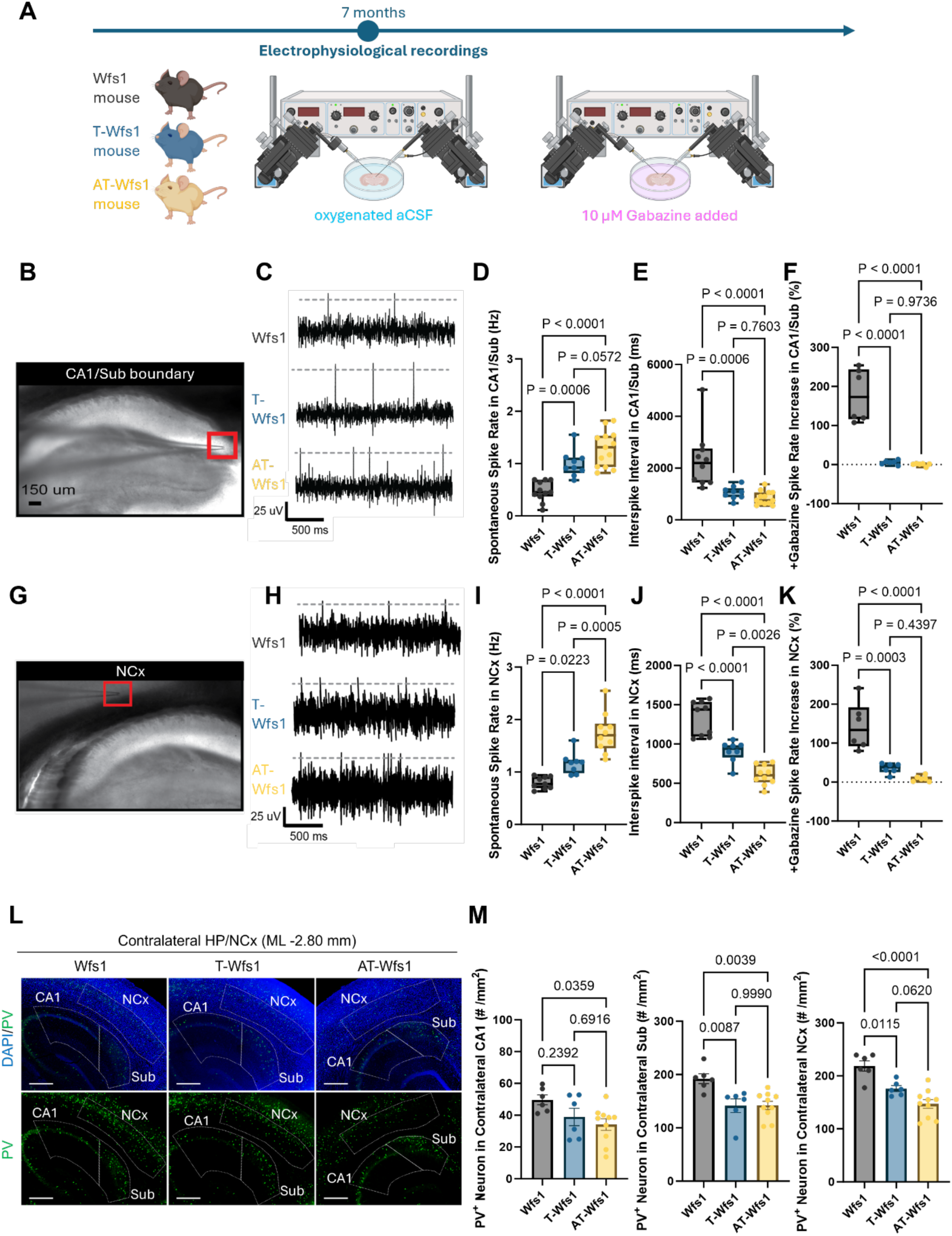
Baseline spontaneous activity of excitatory neurons and distribution of GABAergic neurons. (**A**) The scheme of the experimental design for electrophysiological analysis. Electrophysiological recordings under gabazine-treated and untreated conditions were performed with Wfs1, T-Wfs1, and AT-Wfs1 mice without injection of AAV-FLEX-P301L tau at 7 months of age. (**B**) Recording sites of the CA1/Sub boundary (top) and NCx (bottom). (**C**) Representative traces of spontaneous spiking activity of excitatory neurons in the CA1/Sub boundary and NCx. (**D, I**) Average spontaneous spike rate in each mouse model (n=5 mice per line; n=15 mice total) in the CA1/Sub boundary and NCx. (**E, J**) Average spontaneous interspike interval of each mouse model (n=5 mice per line; n=15 mice total) in the CA1/Sub boundary and NCx. (**F, K**) Average spontaneous spike rate increase (as a percent) of each mouse model (n=2 mice per line; n=6 mice total) in the CA1/Sub boundary and NCx after bath exposure of 10 μM gabazine, a competitive antagonist of GABA_A_ receptors. Means ± SEM. Statistical analyses were performed by one-way ANOVA with Post Hoc Tukey’s multiple comparison test. (**L**) Representative immunofluorescence images of PV (green) and DAPI (blue) in the hippocampal and NCx region of Wfs1, T-Wfs1, and AT-Wfs1 mice. Scale bar = 100 mm. (**M**) Quantification of PV^+^ neurons in the contralateral CA1, Sub, and NCx at ML +2.80 mm. Means ± SEM. Wfs1 n = 6; T-Wfs1 n = 6; AT-Wfs1 n = 10 mice. Statistical analyses were performed by one-way ANOVA with Post Hoc Tukey’s multiple comparison test.

To understand the role of GABAergic signaling in the excitability differences observed in our mouse models, field recordings were conducted in the CA1/Sub boundary and NCx with pharmacological manipulation using the GABA_A_ receptor antagonist gabazine. Following the introduction of gabazine (10 µM) at the CA1/Sub boundary, minimal changes in spontaneous spiking frequency (% change) in T-Wfs1 (4.5 ± 6.9%; n = 5 slices) and AT-Wfs1 (0.48 ± 4.3%; n = 6 slices) groups were observed. In contrast, Wfs1 groups exhibited substantial increases in spiking activity (181 ± 76%; n = 6 slices) (Fig. 5F), suggesting impaired basal GABAergic inhibition of CA1/Sub neurons in the T-Wfs1 (P < 0.0001) and AT-Wfs1 (P < 0.0001) groups compared to Wfs1 controls. In the NCx, distinct changes in spontaneous spiking frequency were observed (% change) after the introduction of gabazine in Wfs1 (150 ± 72%; n = 6 slices), T-Wfs1 (34 ± 15%; n = 5 slices), and AT-Wfs1 (9.7 ± 8.1%; n = 6 slices) mice (Fig. 5K). The significant differences between Wfs1 control and T-Wfs1 (P = 0.0003) or AT-Wfs1 (P < 0.0001) underscore that the combined presence of Tau and Aβ can amplify GABAergic signaling dysfunction more than just Tau alone in the NCx.

### Tau propagation-independent loss of inhibitory neurons in T-Wfs1 and AT-Wfs1 mice

To investigate whether the impaired GABAergic transmission and promoted neuronal hyper excitability were caused by tau propagation-independent loss of inhibitory neurons, count of PV^+^ cells in the contralateral hemisphere of three mouse groups were analyzed. PV^+^ cell counts significantly decreased in the contralateral CA1 (34.09 ± 3.605 cells /mm^2^, P = 0.0359), Sub (142.1 ± 7.813 cells /mm^2^, P = 0.0039) (Fig. 5L-M), and NCx (147.2 ± 8.295 cells /mm^2^, P < 0.0001) of AT-Wfs1 mice (n = 10) (Fig. 6N), and in the contralateral Sub (141.6 ± 12.90 cells /mm^2^, P = 0.0087) and NCx (176.0 ± 5.540 cells /mm^2^, P = 0.0115) of T-Wfs1 mice (n = 10) compared to Wfs1 controls (CA1: 49.68 ± 3.086 cells /mm^2^, Sub: 192.4 ± 9.133 cells /mm^2^, NCx: 218.8 ± 9.738 cells /mm^2^; n = 6) (Fig. 5O). These results indicate that endogenously expressed human *MAPT* and APP^NL-G-F^ synergistically mediate loss of GABAergic inhibitory inhibitory neurons, resulting in the increase of neuronal excitability in these brain regions.

**Fig. 6.**
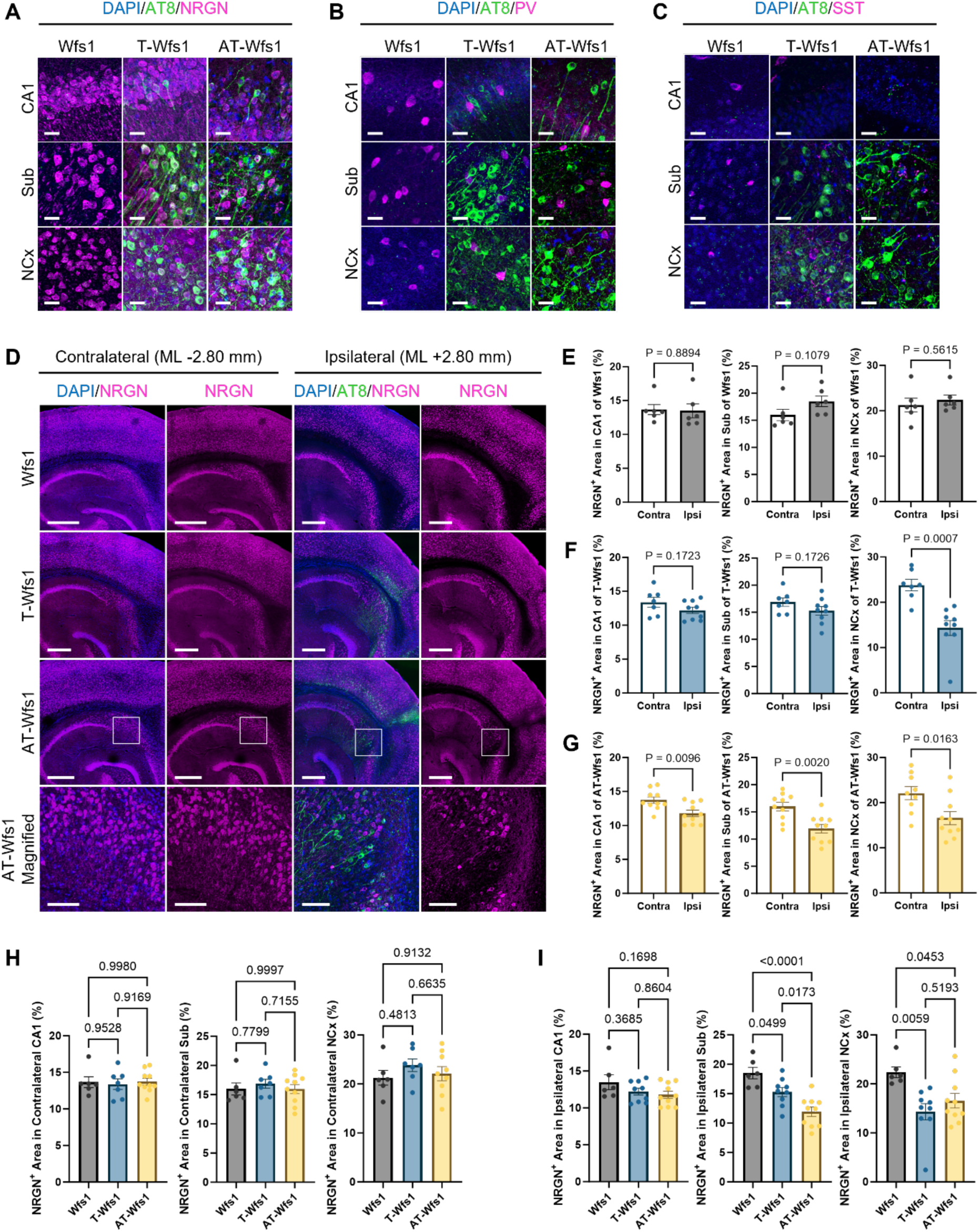
The cell type of the recipient cells targeted of p-Tau propagation. (**A-C**) Representative immunofluorescence image of DAPI (blue), AT8 (green), Neurogranin (NRGN; excitatory neuronal marker, magenta, **A**), Parvalbumin (PV; magenta, inhibitory neuronal marker, **B**), Somatostatin (SST, total Tau, magenta, **C**) in the ipsilateral CA1, Sub, and NCx region at the medial point (ML +2.80 mm, **C**). Scale bar = 100 μm. (**D**) Representative immunofluorescence image of DAPI (blue), AT8 (green), and NRGN (magenta) in the contralateral (left two columns) and the ipsilateral (right two columns) hippocampal/neocortical region at the medial point (ML +2.80 mm). Scale bar = 400 μm or 100 μm in the images in top three row or in the magnified images. (**E-G**) Quantification of the NRGN^+^ area occupancy in the contralateral and the ipsilateral in CA1, Sub, and NCx at ML-2.80 mm and +2.80 mm. Means ± SEM, Wfs1 n = 6 (**E**); T-Wfs1 n = 10 (**F**); AT-Wfs1 n = 10 (**G**) mice. Statistical analyses were performed by Unpaired Student’s t-test. (**H**, **I**) Quantification of the NRGN^+^ area occupancy in the contralateral (**H**) and ipsilateral CA1, Sub, and NCx at ML-2.80 mm or +2.80 mm (**I**). Means ± SEM, Wfs1 n = 6; T-Wfs1 n = 10; AT-Wfs1 n = 10 mice. Statistical analyses were performed by one-way ANOVA with Post Hoc Tukey’s multiple comparison test.

### Excitatory neurons were more vulnerable to propagation and accumulation of p-tau than inhibitory neurons

To characterize neuronal vulnerability for p-tau propagation and accumulation, cell type specific markers colocalized with AT8^+^ cells were profiled in the brains three months post injection of AAV2/6-FLEX-Tau^P301L^ in MECII by co-immunostaining AT8 with anti-neurogranin (NRGN) antibody as an excitatory neuronal marker, and anti-parvalbumin (PV) and somatostatin (SST) antibodies as inhibitory neuronal markers. In all three groups, AT8^+^ cells were most frequently colocalized with NRGN^+^ excitatory neuron. A total of 553 out of 1596 AT8^+^ cells (34.6%) were most frequently colocalized with NRGN^+^ excitatory neuron, while 1 out of 2529 AT8^+^ cells (0.0395%) or 20 out of 1902 AT8^+^ cells (1.05%) of AT8^+^ cells were colocalized with PV^+^ or SST^+^ inhibitory neurons respectively (Table 1, Fig. 6A–C). Furthermore, the NRGN^+^ area occupancy significantly decreased in ipsilateral hemi-brains compared with contralateral hemi-brains in NCx of T-Wfs1 mice (Fig. 6D, F) and in CA1, Sub, and NCx of AT-Wfs1 mice (Fig. 6D, G), while there was no difference of NRGN^+^ area occupancy between ipsilateral and contralateral hemi-brains of Wfs1 mice (Fig. 6D, E). Additionally, there was no differences of the NRGN^+^ area occupancy in contralateral hemisphere among three groups (Fig. 6D, H), while decrease of the NRGN^+^ area occupancy in ipsilateral hemisphere of T-Wfs1 and AT-Wfs1 mice compared with Wfs1 mice was observed (Fig. 6D, I). The spatial correlation between NRGN^+^ excitatory neuronal loss and AT8^+^ cells in these regions underscores a direct link between excitatory neuron degeneration and Tau spread. These results suggested that excitatory neurons in T-Wfs1 and AT-Wfs1 mice were more vulnerable to tau propagation than those in Wfs1 mice.

**Table 1.** The colocalization of co-localization of AT8 positive cells and specific cell type markers positive cells. Count (#) and ratio (%) of co-localization of AT8^+^ cells and specific cell type markers positive cells: NRGN (left column), PV (middle column), and SST (right column), in the ipsilateral CA1, Sub, and NCx region of Wfs1, T-Wfs1, and AT-Wfs1 mice at the medial point (ML +2.80 mm). Scale bar = 100 μm. Value shows the total number of the cells counted in 3 sections at around ML +2.80 to +2.00 mm from each mouse, Wfs1 n = 6; T-Wfs1 n = 6; AT-Wfs1 n = 5 mice.

|  |  | AT8<sup>+</sup>NRGN<sup>+</sup><br>/AT8<sup>+</sup> |  | AT8<sup>+</sup>PV<sup>+</sup><br>/AT8<sup>+</sup> |  | AT8<sup>+</sup>SST<sup>+</sup><br>/AT8<sup>+</sup> |  |
| --- | --- | --- | --- | --- | --- | --- | --- |
|  |  | # | % | # | % | # | % |
| **Wfs1** | **VC** | 14/14 | 100 | 0/4 | 0 | 0/4 | 0 |
|  | **Sub** | 39/45 | 86.7 | 0/51 | 0 | 1/56 | 1.79 |
|  | **CA1** | 2/2 | 100 | 0/0 | NA | 0/0 | NA |
|  | **Total** | 55/61 | 90.2 | 0/55 | 0.0 | 1/60 | 1.67 |
| **T-Wfs1** | **VC** | 174/581 | 29.9 | 0/560 | 0 | 10/549 | 1.82 |
|  | **Sub** | 140/283 | 49.5 | 0/413 | 0 | 0/372 | 0 |
|  | **CA1** | 28/38 | 73.7 | 1/231 | 0.433 | 0/3 | 0 |
|  | **Total** | 342/902 | 37.9 | 1/1204 | 0.0831 | 10/924 | 1.08 |
| **AT-Wfs1** | **VC** | 114/524 | 21.8 | 0/563 | 0 | 6/636 | 0.943 |
|  | **Sub** | 38/92 | 41.3 | 0/546 | 0 | 3/237 | 1.27 |
|  | **CA1** | 4/17 | 23.5 | 0/161 | 0 | 0/45 | 0 |
|  | **Total** | 156/633 | 24.6 | 0/1270 | 0 | 9/918 | 0.980 |
| **Total** |  | 553/1596 | 34.6 | 1/2529 | 0.0395 | 20/1902 | 1.05 |

## Discussion

This study offers novel mechanistic insights into circuit-specific and cell-type-dependent propagation of Tau pathology in genetically modified mouse models of AD. By combining viral-mediated Tau expression and detailed neuroanatomical mapping, the influence of genetic background, neuronal excitability, and projection architecture on Tau pathology was established: mirroring key aspects of early Braak staging in AD.

### Genetic context dictates tau propagation from MECII

This study demonstrates that tau propagation from the MECII to the neocortical region through the hippocampal region depends on both human tau expression and amyloidogenic context. MECII-injected human tau propagation was restricted from MECII to proximal CA1 and distal Sub in Wfs1 mice, whereas tau propagated to the NCx region adjacent to Sub region visual cortex as well as HP region in T-Wfs1 and AT-Wfs1 mice. Furthermore, while the tau projection to proximal CA1 remained intact in T-Wfs1 mice, AT-Wfs1 mice showed tau projection to CA1/Sub boundary (distal CA1 and proximal Sub) and no detectable tau in proximal CA1, which may be attributable to reduced dendritic integrity in the SR region of proximal CA1. This observation aligns with prior reports suggesting that early tau pathology appears near the CA1/CA2 boundary in PART, which is defined as the presence of Alzheimer-type neurofibrillary pathology in the medial temporal lobe and other structures in the absence of significant Aβ plaque deposition, while the CA1/Sub boundary shows increased susceptibility to tau pathology in association with Aβ plaque deposition in AD (Walker, Fudym et al. 2021, Walker, Richardson et al. 2021). Additionally, CA1/Sub region also have been known that the most vulnerable to Aβ pathogenesis in hippocampal region as Aβ plaque has already been observed in the Thal phase 1 in AD brains (Thal, Rub et al. 2002). Therefore, our newly developed T-Wfs1 and AT-Wfs1 mouse models may serve as useful platforms to model tau pathology formation in PART and AD, respectively, and suggest that Aβ plaque pathology contributes to differences in tau propagation routes between PART- and AD-like conditions.

The differential burden of AT8 and Alz50 signal in T-Wfs1 versus AT-Wfs1 mice underscores the synergistic interaction between tau and Aβ, a relationship also supported by human imaging studies that illustrate Aβ accumulation precedes and predicts the onset of Tau pathology in connected cortical regions (Jack, Knopman et al. 2013, Hanseeuw, Betensky et al. 2019). Our finding that AT-Wfs1 mice exhibit more misfolded tau burden along with neuronal cell loss than Wfs1 or T-Wfs1 mice supports the previous findings of enhanced tau toxicity by Aβ deposition in human brains (Ittner and Götz 2011, Bloom 2014). Aβ has been reported to activate representative tau kinases such as glycogen synthase kinase-3β, cyclin dependent kinase 5 and most recently tau-tubulin kinase 1, promoting tau hyperphosphorylation and axonal degeneration (Noble, Olm et al. 2003, Ikezu, Ingraham Dixie et al. 2020), and may facilitate tau misfolding through oxidative stress (Abramov, Potapova et al. 2020). This finding suggests that Aβ not only enhances tau pathology but may also facilitate conformational changes required for their toxicity and neurotoxicity through mitochondrial damage.

### Circuit-specific transmission via anatomical connectivity

Our tracing experiments reveal that the Sub/CA1 boundary mediates tau propagation to NCx areas such as the V1 region. Injection of AAV-FLEX-Tau^P301L^ into the Sub, but not proximal CA1, produced tau accumulation in both Sub/CA1 boundary and NCx region, indicating projection-specific model of tau spread, where regional vulnerability is dictated by axonal projection architecture and synaptic integration. This selective spread mirrors early Braak stages, where tau pathology spreads from EC to HP and then to higher-order association cortices (Braak and Braak 1991), suggesting that vulnerability to tau pathology is structured by underlying connectivity. Ultimately, these results support the growing consensus that tau pathology is dependent upon specific neural circuitry and propagates along anatomically connected pathways (de Calignon, Polydoro et al. 2012, Liu, Drouet et al. 2012, Vogel, Iturria-Medina et al. 2020).

### Network dysfunction reflects impaired inhibitory tone and hyperexcitability

A major mechanistic insight from our study is the role of neuronal excitability in modulating tau spread. Field recordings revealed increased spontaneous spiking and reduced interspike intervals in both T-Wfs1 and AT-Wfs1 mice relative to Wfs1 controls, particularly at the CA1/Sub boundary and in the NCx. These findings are consistent with electrophysiological studies showing hyperactivity in early AD models and patients (Busche and Konnerth 2015). Notably, pharmacological blockade of GABA_A_ receptors with gabazine revealed a profound disruption of inhibitory control in both tau and Aβ models, with the most pronounced deficits observed in AT-Wfs1 mice indicating a significant loss of GABAergic inhibitory network in AT-Wfs1 mice. This result aligns with prior studies as evidence has shown that tau pathology can disrupt GABAergic inhibitory neuron function and synaptic inhibition (Verret, Mann et al. 2012), and that this imbalance of excitatory-inhibitory neurotransmission can lead to network instability, epileptiform activity, and cognitive impairment (Palop and Mucke 2010, Ranasinghe, Kudo et al. 2025). Ultimately, our results support a model in which tau-induced dysfunction of GABAergic inhibition contributes to hyperexcitability, which in turn may amplify tau propagation, creating a feed-forward loop of circuit deterioration.

### Cell Type-Specific Vulnerability Reflects Intrinsic Susceptibility

We observed a preferential accumulation of p-tau in NRGN^+^ excitatory neurons, with relatively sparse involvement of SST^+^ inhibitory neurons and a near absence in PV^+^ GABAergic inhibitory neurons. This pattern aligns with prior findings from both human and murine studies indicating that excitatory pyramidal neurons are the earliest and most vulnerable to tau pathology (Yoshiyama, Higuchi et al. 2007, Murray, Przybelski et al. 2014). The selective vulnerability of excitatory neurons may stem from their high synaptic activity, susceptibility to calcium dysregulation and tau accumulation, and expression of proteins that mediate tau internalization or seeding (Zempel and Mandelkow 2015). However, clear loss of GABAergic input in T-Wfs1 and AT-Wfs1 mice indicates that although the number of PV neurons are preserved, their functions are significantly impaired in the context of GABAergic transmission.

## Conclusion

Our study establishes a comprehensive mechanistic framework for early tau pathology progression in AD by demonstrating how tau propagation is modulated by genetic context, projection-specific neuronal connectivity, and activity-dependent processes. Using the Wfs1-based models expressing human tau and Aβ, we show that amyloid pathology amplifies the spread of tau and its pathological transformation into phosphorylated and misfolded isoforms. We also identify the Sub region as a critical hub to enable cortical tau dissemination, highlighting the importance of distinct anatomical pathways in shaping pathological spread relevant to the limbic stage of AD pathology development. Importantly, our findings reveal that excitatory neurons are selectively vulnerable to tau accumulation and degeneration, while GABAergic signaling dysfunction contributes to network hyperexcitability. likely facilitating further tau spread. In conclusion, these findings illuminate multiple converging mechanisms, genetic, anatomical, cellular, and physiological, that synergistically drive early tau pathology in AD. Ultimately these results point towards circuit-specific and excitability-modulating strategies as potential therapeutic avenues to halt disease progression.

## Acknowledgements

We would like to thank S. Tonegawa for providing the Wfs1-Cre mice, P. Davies, for the gift of the Alz50 monoclonal antibodies, and members of the Laboratory of Molecular Neurotherapeutics for scientific suggestions and technical assistance.

## Author contributions

T.I. conceptualized the study, performed vector design, construction, and validation. A.R.R. performed animal injections, immunohistochemistry, confocal microscopy. A.H.-T. organized immunohistochemistry study and performed immunohistochemistry, confocal microscopy, and data analysis. B.J.G. conceptualized and organized electrophysiology studies and performed electrophysiology analysis. J.E., S.R., N.B.R., and A.L.-P. performed immunohistochemistry, confocal microscopy, and data analysis. A.R.R., A.H.-T., B.J.G., J.E., N.B.R., A.L.-P., S.I., and T.I. wrote and edited the manuscript. We would like to thank BioRender for providing the platform to create the figures in this manuscript.

