## Supplementary figures for "Humanized tau and amyloid-β deposition accelerate tau propagation, neuronal cell loss and neurophysiological dysfunction in novel mouse models of primary age-related tauopathy and Alzheimer’s disease"

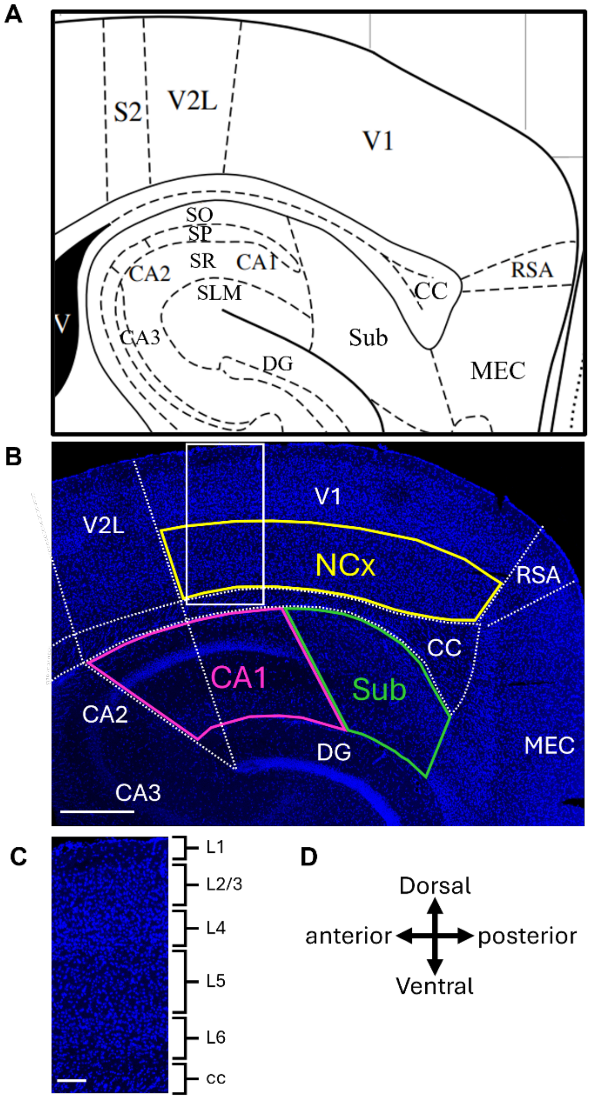


**Fig. S1. The segment three ROIs for the analysis of signal positive area occupancy.**

(**A**) Anatomical map image of mouse sagittal section at ML 2.76 mm. (**B**) Representative immunofluorescence image of DAPI (blue) images in hippocampal/neocortical region in the sagittal section. In the analysis of signal-positive area occupancy, three ROIs for CA1 (magenta-outlined region), subiculum (Sub, green-outlined region), and neocortex (NCx, yellow-outlined region) were segmented. Scale bar = 400 µm. (**C**) Magnified imaged of boxed area in NCx. Scale bar = 100 µm. (**D**) Direction arrows in anatomical map. V1: primary visual cortex, V2L: secondary visual cortex, S2: secondary somatosensory cortex lateral area, RSA; retrosplenial agranular cortex, MEC: medial entorhinal cortex, Sub: subiculum, CA1-3: field CA1-3 of hippocampus, SO: stratum oriens, SP: stratum pyramidal, SR: stratum radiatum, SLM: stratum lacunosum-moleculare, DG: dentate gyrus.

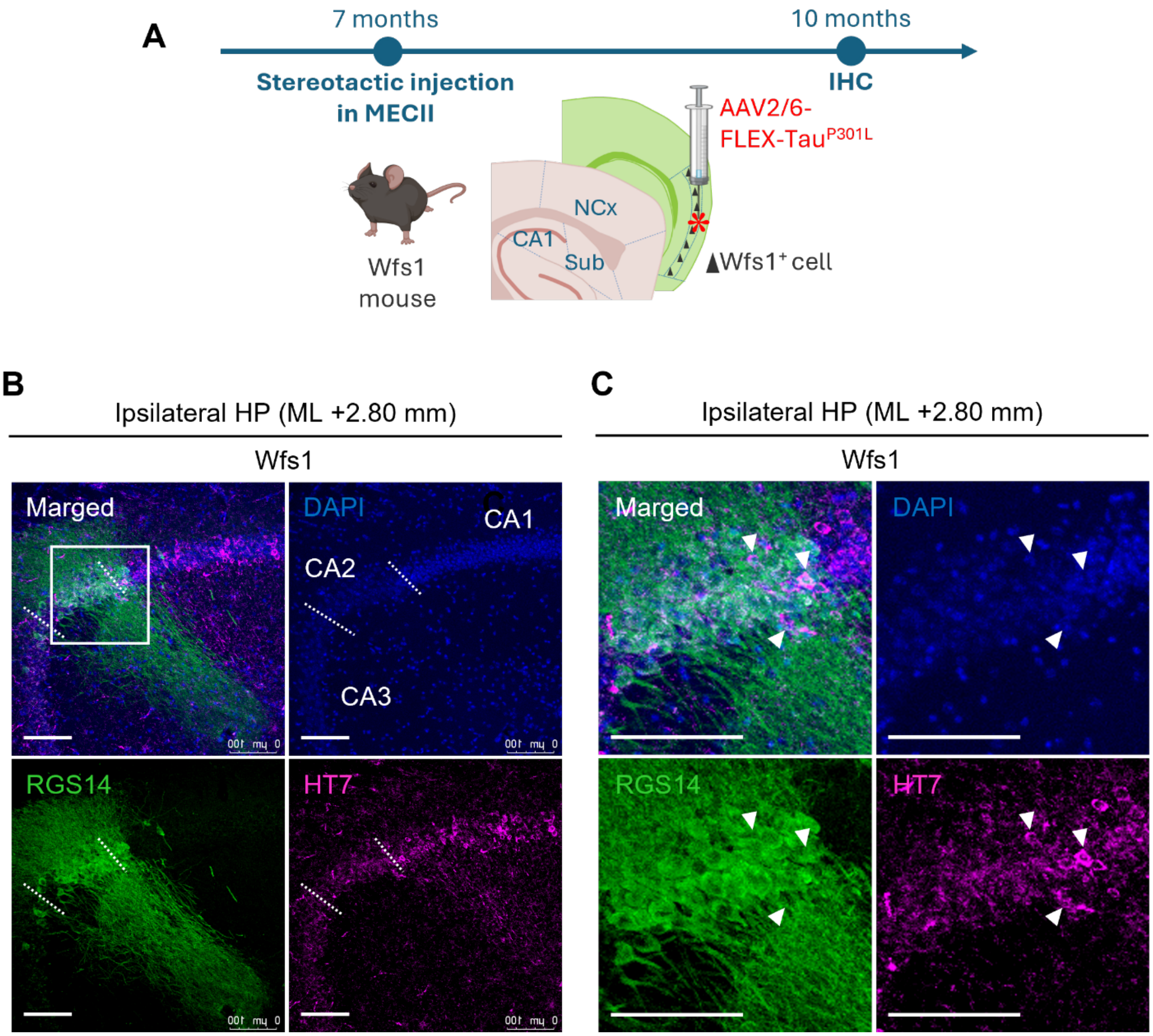


**Fig. S2. The tau propagation into CA1 pyramidal cells around the CA1/CA2 boundary of Wfs1 mouse.**

(**A**) Experimental design schematic for AAV2/6-FLEX-Tau$^{P301L}$ injection in Wfs1 mouse. Animals were injected with AAV2/6-FLEX-Tau$^{P301L}$ in the MECII (AP -4.85 mm, ML +3.45 mm, DV -3.30 mm) at 7 months of age. Mice are euthanized 3 months after the injection for immunohistochemistry. (**B**) Representative immunofluorescence image of DAPI (blue), RGS14 (CA2 neuron, green) and HT7 (total Tau, magenta) in the ipsilateral CA1/CA2 boundary of Wfs1 mouse at the medial point (ML +2.80 mm). Dashed lines indicate the CA1/CA2 and CA2/CA3 boundaries defined based on the DAPI image. Scale bar = 100 μm. (**C**) Magnified image of the boxed area. Scale bar = 100 μm Arrowheads point to HT7$^{+}$ somata at the CA1/CA2 border. The HT7$^{+}$ signal exhibited a soma size comparable to that of CA1 pyramidal neurons and did not co-localize with RGS14; therefore, these findings indicate that tau propagates to proximal CA1 pyramidal neurons.

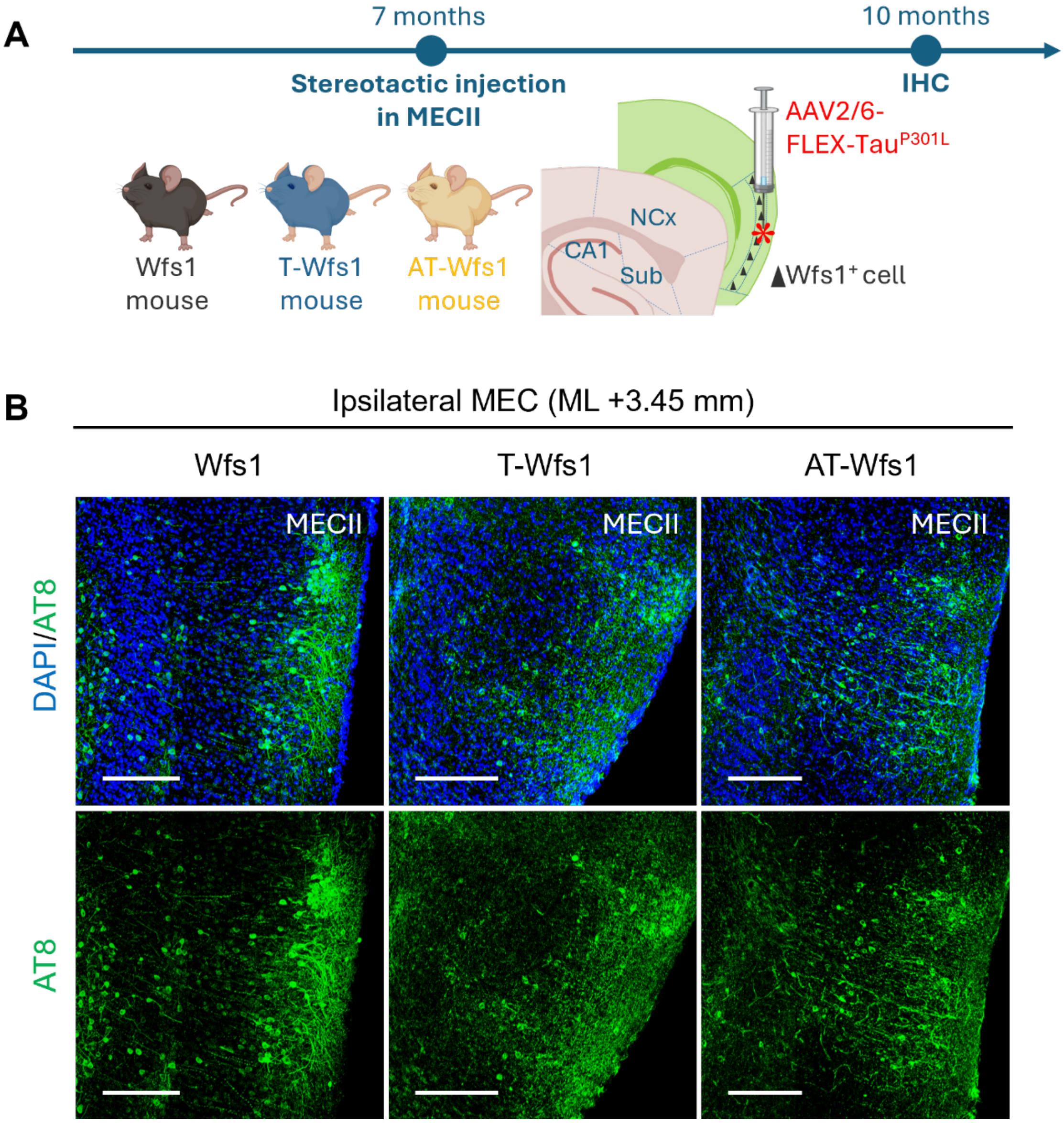


**Fig. S3. The accumulation of p-tau in the MECII in Wfsq, T-Wfs1, AT-Wfs1 mice.**

(**A**) Experimental design schematic for AAV2/6-FLEX-Tau$^{P301L}$ injection in Wfs1, T-Wfs1, and AT-Wfs1 mice. Animals were injected with AAV2/6-FLEX-Tau$^{P301L}$ in the MECII (AP -4.85 mm, ML +3.45 mm, DV -3.30 mm) at 7 months of age. Mice are euthanized 3 months after the injection for immunohistochemistry. (**A**) Representative immunofluorescence image of DAPI (blue) and AT8 (pS$^{202}$/pT$^{205}$ Tau, green) in the MECII region of three groups at the injection point (ML +3.45 mm). Scale bar = 200 µm.

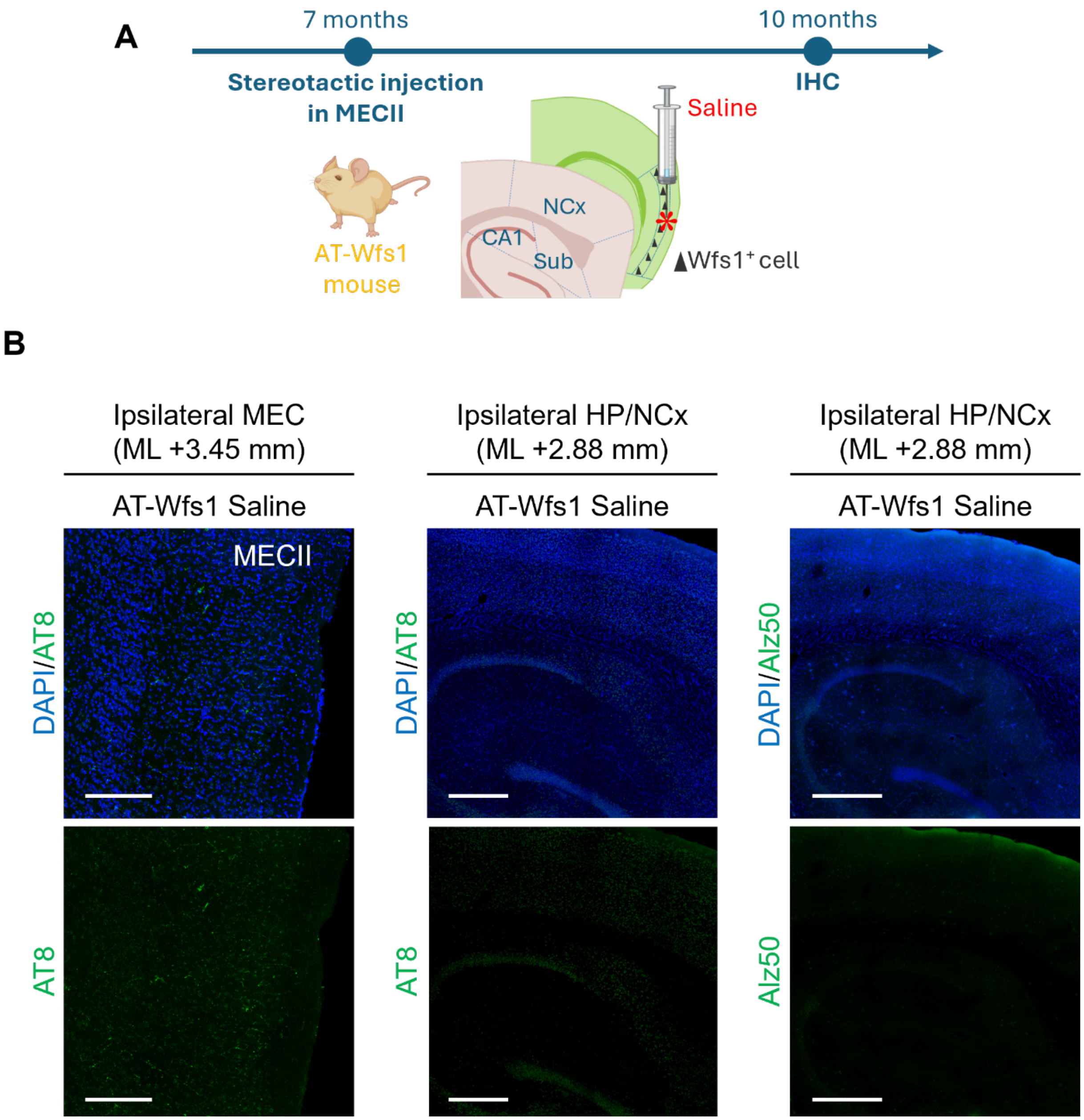


**Fig. S4. No p-tau signal detected in the ipsilateral hemisphere of AT-Wfs1 mouse injected with sline.**

(**A**) Experimental design schematic for AAV2/6-FLEX-Tau$^{P301L}$ injection in AT-Wfs1 mice. Animals were injected with saline in the MECII (AP -4.85 mm, ML +3.45 mm, DV –3.30 mm) at 7 months of age. Mice are euthanized 3 months after the injection for immunohistochemistry. (**B**, **C**) Representative immunofluorescence image of DAPI (blue), AT8 (green) in the ipsilateral MEC region at at the injection point (ML +3.45 mm, Scale bar = 200 µm, **B**) and hippocampal/neocortical region of AT-Wfs1 mouse at the medial point (ML +2.80 mm, Scale bar = 200 µm, **C**). (**D**) Representative immunofluorescence image of Alz50 (misfolded Tau, green) in the ipsilateral hippocampal/neocortical region of AT-Wfs1 mouse at the medial point (ML +2.80 mm). Scale bar = 400 µm.

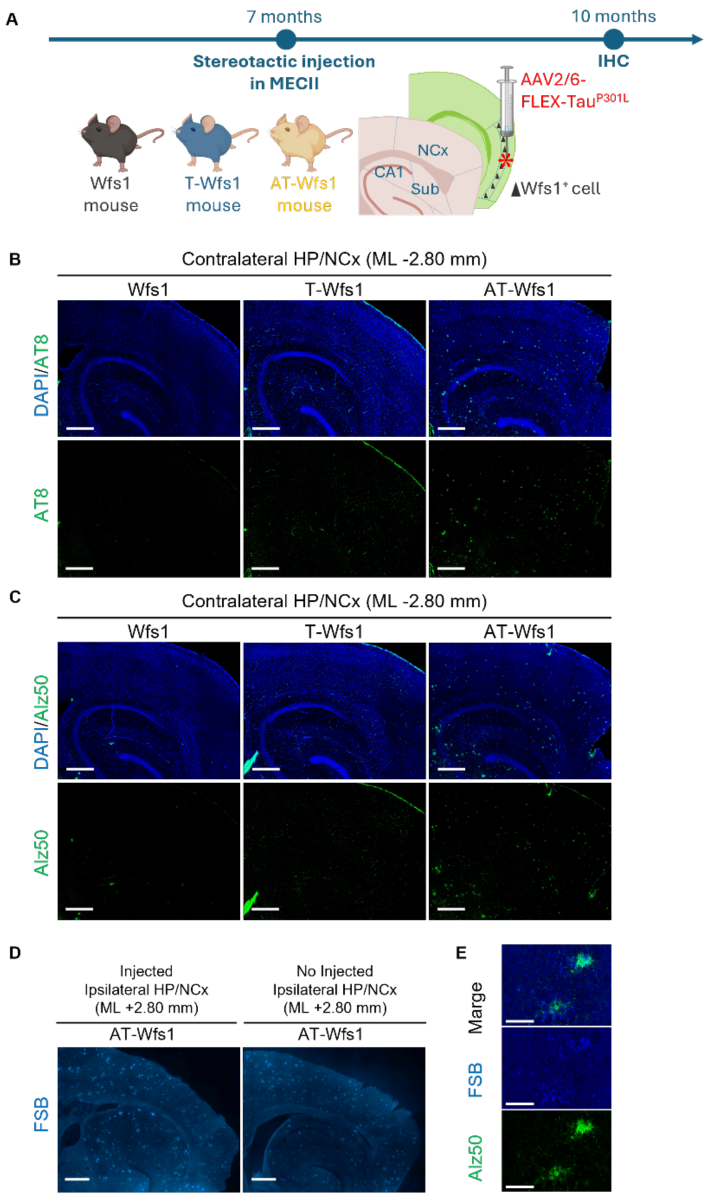


**Fig. S5. The distribution of propagated Tau into the contralateral hemisphere and the deposition of Aβ plaque.**

(**A**) Experimental design schematic for AAV2/6-FLEX-Tau$^{P301L}$ injection in Wfs1, T-Wfs1, and AT-Wfs1 mice. Animals were injected with AAV2/6-FLEX-Tau$^{P301L}$ in the MECII (AP –4.85 mm, ML +3.45 mm, DV –3.30 mm) at 7 months of age. Mice are euthanized 3 months after the injection for immunohistochemistry. (**B**, **C**) Representative immunofluorescence image of DAPI (blue), AT8 (green, **B**) and Alz50 (green, **C**) in the contralateral hippocampal/neocortical region of three groups at the medial point (ML +2.80 mm). Scale bar = 400 µm. (**D**) Representative immunofluorescence image of FSB (Aβ plaque, blue) in the ipsilateral hippocampal/neocortical region of T-Wfs1 and AT-Wfs1 mice with or without of AAV2/6-FLEX-Tau$^{P301L}$ injection at the medial point (ML +2.80 mm). Scale bar = 400 µm. (**E**) Representative immunofluorescence image of FSB (Aβ plaque, blue) and Alz50 (misfolded Tau, green) in AT-Wfs1 mice with AAV2/6-FLEX-Tau$^{P301L}$ injection. Scale bar = 50 µm.

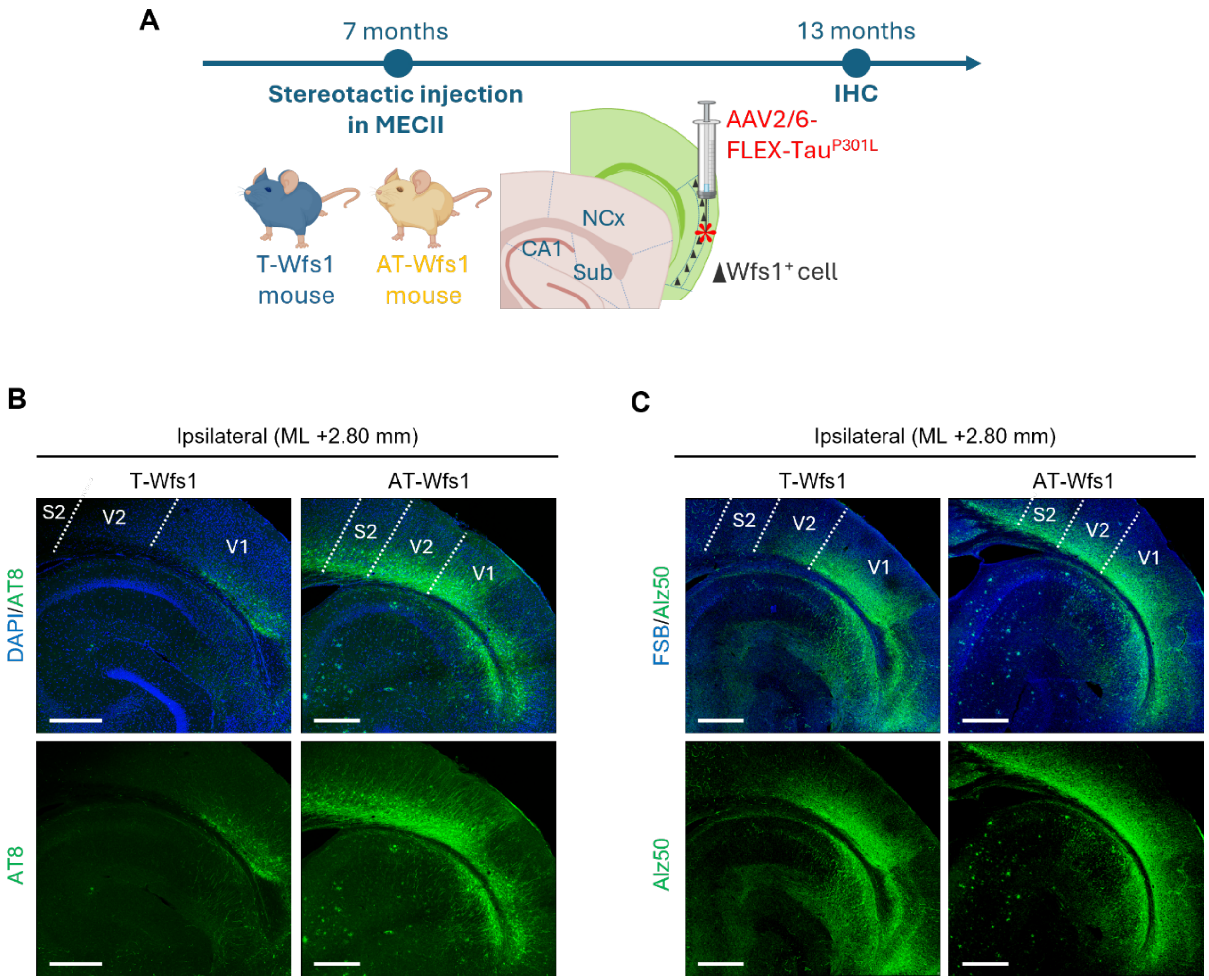


**Fig. S6. Expansion of tau propagation regions with prolonged incubation time after AAV2/6-FLEX-Tau$^{P301L}$ injection.**

(**A**) Experimental design schematic for AAV2/6-FLEX-Tau$^{P301L}$ injection in T-Wfs1 and AT-Wfs1 mice. Animals were injected with AAV2/6-FLEX-Tau$^{P301L}$ in medial entorhinal cortex layer 2 (MECII; AP –4.85 mm, ML +3.45 mm, DV 3.30 mm) at 7 months of age. Mice are euthanized 6 months after the injection for immunohistochemistry. (**B**) Representative immunofluorescence image of DAPI (blue) and AT8 (green) in the ipsilateral hippocampal/neocortical region of T-Wfs1 and AT-Wfs1 mice at the medial point (ML +2.80 mm). Scale bar = 400 µm. (**C**) Representative immunofluorescence image of FSB (Aβ plaque, blue) and Alz50 (misfolded Tau, green) in the ipsilateral hippocampal/neocortical region of T-Wfs1 and AT-Wfs1 mice at the medial point (ML +2.80 mm). Scale bar = 400 µm

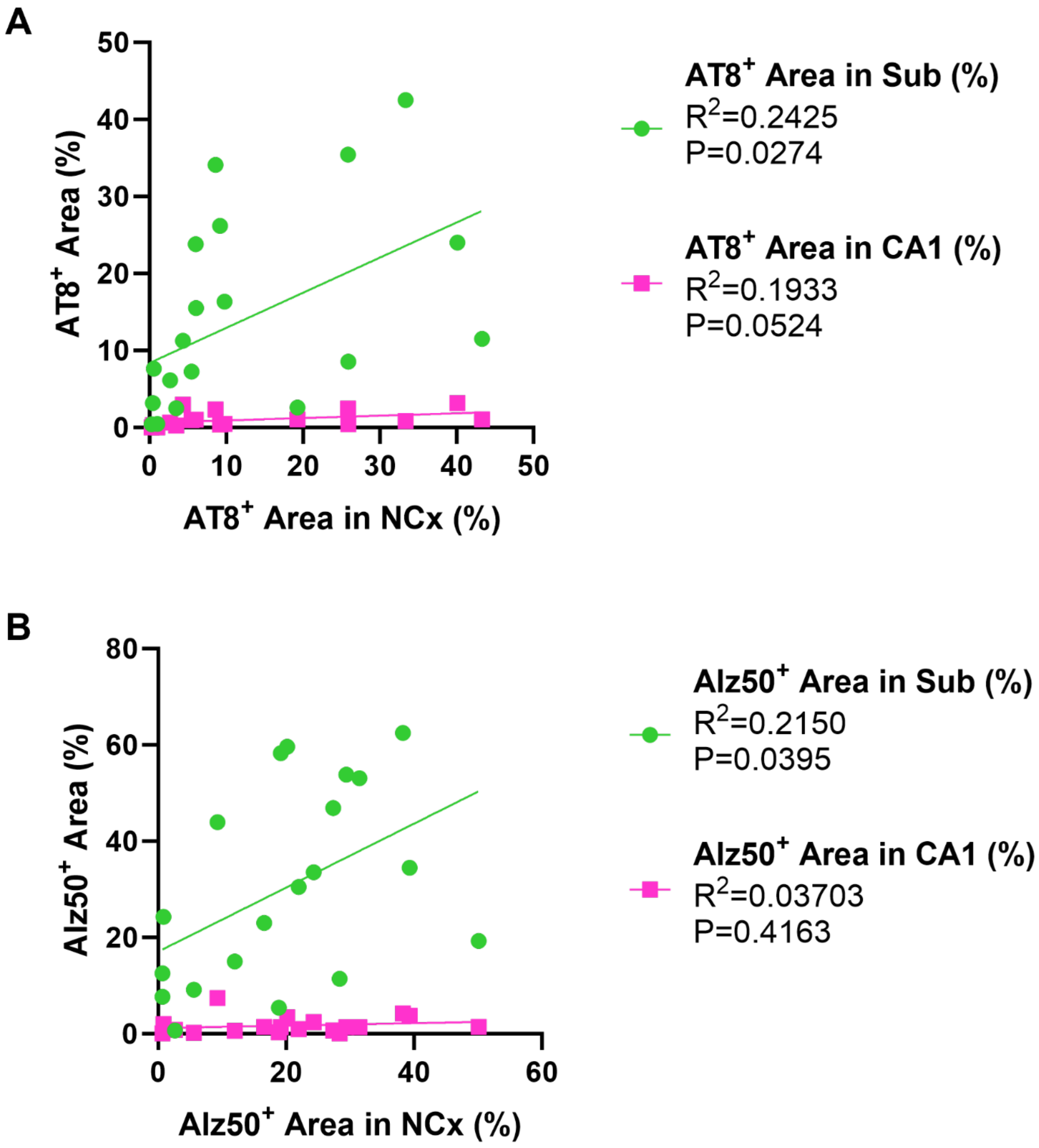


**Fig. S7. Correlation of the area occupancy of p-Tau and misfolded Tau between CA1 and NCx or between Sub and NCx.**

(**A**, **B**) Scatter plot showing the area occupancy of $AT8^+$ (**A**) or $Alz50^+$ (**B**) in NCx (x-axis) versus in CA1 or in Sub (magenta or green, y-axis). Each dot represents one sample (T-Wfs1 n = 10; AT-Wfs1 n = 10 mice). The solid line indicates the best-fit linear regression (least squares). The Pearson's correlation coefficient of determination ($R^2$) and P value were calculated in GraphPad Prism using the Correlation Matrix, with two-tailed testing.

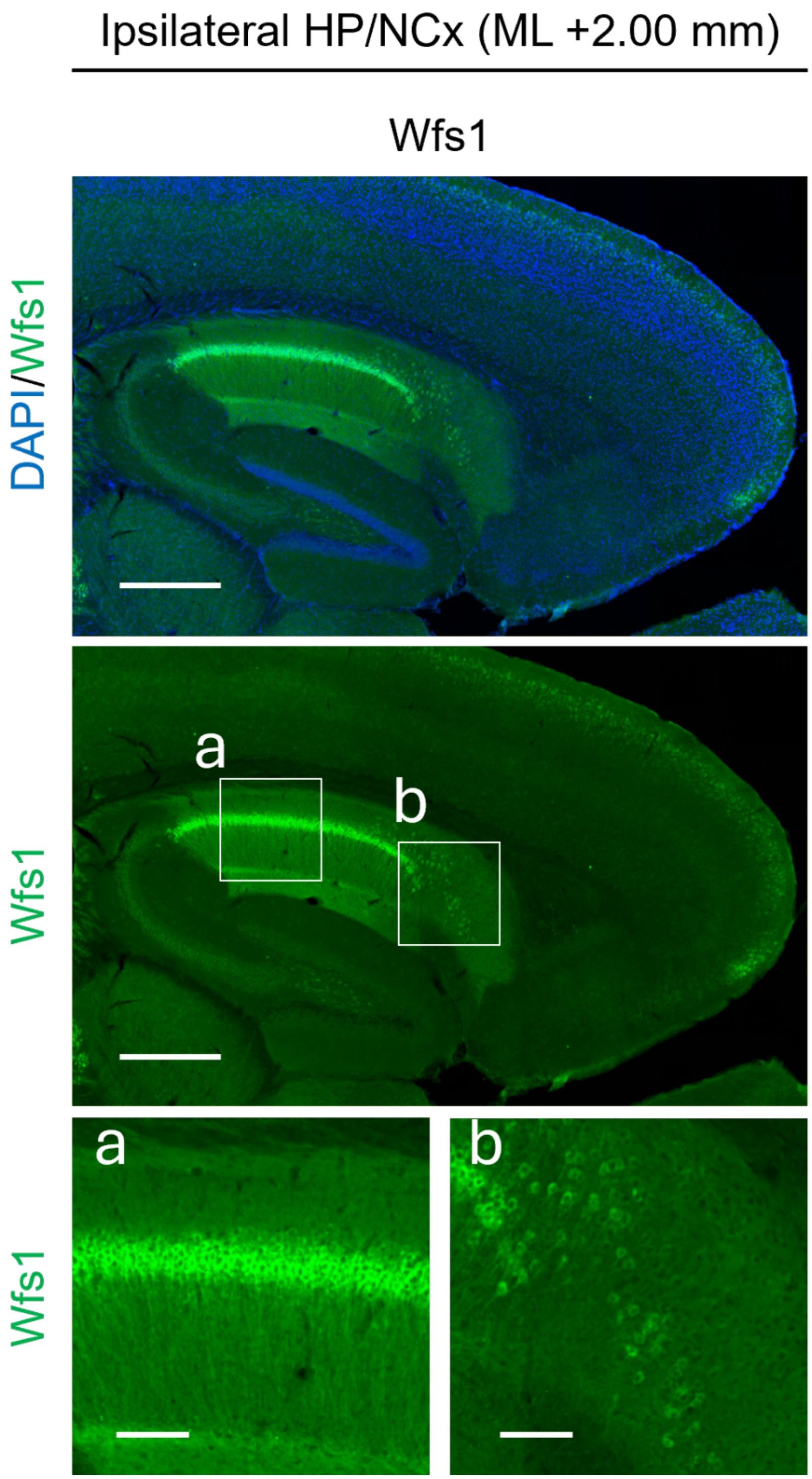


**Fig. S8. The localization of Wfs1$^{+}$ cells in CA1 and Sub.**

Representative immunofluorescence image of DAPI (blue) and Wfs1 (green) in the ipsilateral hippocampal/neocortical region of Wfs1 mice at the medial point (ML +2.00 mm). Scale bar = 500 µm. (**a, b**) Magnified image of the boxed area in CA1 (**a**) and Sub (**b**). Scale bar = 100 µm

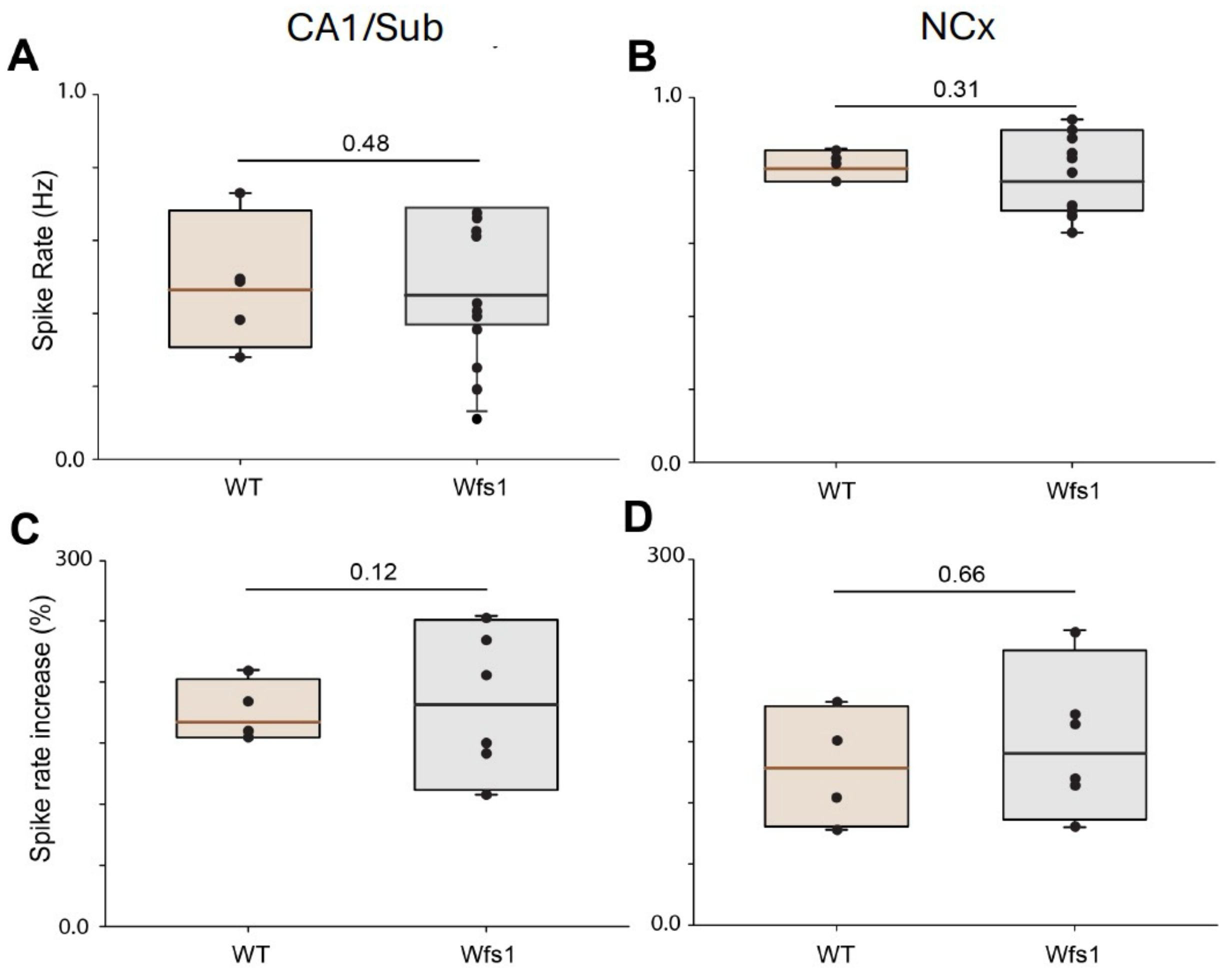


**Fig. S9. Baseline spontaneous activity of excitatory neurons in the CA1/Sub boundary and NCx of WT and Wfs1 mice.**

(**A**-**D**) Average spontaneous spike rate of each control line (WT: n = 2 mice, Wfs1: n = 5 mice) in the CA1/Sub boundary (WT n = 5 slices, Wfs1 n = 11 slices) (**A**) and VC region (WT n = 4 slices, Wfs1 n = 9 slices) (**B**). Average spontaneous spike rate increase (as a percent) of each mouse model (n = 2 mice per line) in the CA1/Sub boundary (WT n = 4 slices, Wfs1 n = 6 slices) (**C**) and VC region (WT n = 4 slices, Wfs1 n = 6 slices) (**D**) after bath exposure of 10 μM gabazine (a specific antagonist of $GABA_A$ receptors). Data represent Means ± SEM. Statistical analyses were performed by unpaired Student's t-test.
